# Basal MET Phosphorylation is an Indicator of Hepatocyte Dysregulation in Liver Disease

**DOI:** 10.1101/2023.07.04.547655

**Authors:** Sebastian Burbano de Lara, Svenja Kemmer, Ina Biermayer, Svenja Feiler, Artyom Vlasov, Lorenza A. D’Alessandro, Barbara Helm, Yannik Dieter, Ahmed Ghallab, Jan G. Hengstler, Katrin Hoffmann, Marcel Schilling, Jens Timmer, Ursula Klingmüller

**Affiliations:** Division Systems Biology of Signal Transduction, German Cancer Research Center (DKFZ), Heidelberg, Germany; Institute of Physics, University of Freiburg, Freiburg, Germany; FDM - Freiburg Center for Data Analysis and Modeling, University of Freiburg, Freiburg, Germany; Liver Systems Medicine against Cancer (LiSyM-Krebs), Germany; Department of General, Visceral and Transplant Surgery, Heidelberg University, Heidelberg, Germany; Systems Toxicology, Leibniz Research Center for Working Environment and Human Factors, Technical University Dortmund, Germany; Department of Forensic Medicine and Toxicology, Faculty of Veterinary Medicine, South Valley University, Qena 83523, Egypt; Signalling Research Centres BIOSS and CIBSS, University of Freiburg, Freiburg, Germany

**Keywords:** Western diet, dynamic pathway modeling, hepatocyte dysregulation, fatty liver disease, HGF signal transduction

## Abstract

Chronic liver diseases are worldwide on the rise. Due to the rapidly increasing incidence, in particular in Western countries, Non-alcoholic fatty liver disease (NAFLD) is gaining importance as the disease can develop into hepatocellular carcinoma. Lipid accumulation in hepatocytes has been identified as the characteristic structural change in NAFLD development, but molecular mechanisms responsible for disease progression remained unresolved. Here, we uncover in primary hepatocytes from a preclinical model fed with a Western diet (WD) a strong downregulation of the PI3K-AKT pathway and an upregulation of the MAPK pathway. Dynamic pathway modeling of hepatocyte growth factor (HGF) signal transduction combined with global proteomics identifies that an elevated basal MET phosphorylation rate is the main driver of altered signaling leading to increased proliferation of WD-hepatocytes. Model-adaptation to patient-derived hepatocytes reveal patient-specific variability in basal MET phosphorylation, which correlates with patient outcome after liver surgery. Thus, dysregulated basal MET phosphorylation could be an indicator for the health status of the liver and thereby inform on the risk of a patient to suffer from liver failure after surgery.

## Introduction

Chronic liver disease is a frequent pathology with increasing mortality rates. Hepatocytes, the most abundant cell type in the liver, play a central role in metabolism and for example store and degrade glycogen to ensure a constant supply of glucose in the blood. High caloric intake can disrupt this process and result in fatty liver disease (Riazi *et al*, 2022), a metabolic disorder characterized by the accumulation of lipid droplets in hepatocytes. If no secondary causes such as alcohol abuse, metabolic syndromes, or medication are identified, fatty liver disease is categorized as Non-alcoholic fatty liver disease (NAFLD). In the past decades the incidence of NAFLD has steadily increased. If sustained for a long period of time, NAFLD can develop into non-alcoholic steatohepatitis (NASH), fibrosis, cirrhosis and even hepatocellular carcinoma (Huang *et al*, 2021). Therefore, an understanding of the underlying mechanisms of disease development and progression is pivotal. Metabolic changes in different liver disease stages have been extensively characterized (Jia *et al*, 2014; Puri *et al*, 2007), whereas alterations in information processing of hepatocytes driven by high caloric diets have not yet been investigated in depth. Previous observations (Oe *et al*, 2005; Paranjpe *et al*, 2016; Tekkesin *et al*, 2011) suggest that hepatocyte growth factor (HGF) might play a central role in hepatic pathologies. HGF binds to the receptor tyrosine kinase MET on hepatocytes and triggers the activation of proliferative signal transduction by the mitogen activated kinase pathway (MAPK) and the phosphoinositide 3 kinase (PI3K)/AKT pathway. Through the negative feedback loop between S6K (MAPK) and IRS1 (PI3K/AKT), MET stimulation can affect the metabolic functions of the liver during regeneration (Hall *et al*, 2021). Furthermore, we utilized dynamic pathway modelling to disentangle the cross talk of the MAPK and PI3K/AKT pathways in these cells (D’Alessandro *et al*, 2015) and showed that both, ERK phosphorylation and PI3K activation, are required for proliferation of primary mouse hepatocytes (Mueller *et al*, 2015). As most liver pathologies are driven by increasing damage of the liver, consideration of HGF induced signal transduction in hepatocytes, which is essential for liver regeneration, could be informative to predict patient outcome upon liver surgery. Currently, only postoperative descriptive scores such as the Clavien Dindo score (Dindo *et al*, 2004) or the comprehensive complication index (Slankamenac *et al*, 2013) are documented and so far no correlations with preoperative metrices could be established. The implementation of complex patient datasets for clinical decisions remains challenging and requires the development of comprehensive tools. A suitable approach could be the use of mechanism-based dynamic pathway models to integrate and exploit complex datasets (D’Alessandro *et al*, 2022; Kok *et al*, 2020; Oppelt *et al*, 2018) in order to guide clinical decisions.

In this work, we utilized the Western diet (WD) mouse as a preclinical model to study alterations in HGF signal transduction occurring in liver disease. Data generated from primary murine hepatocytes of healthy and WD mice and from patient-derived primary human hepatocytes were used to calibrate a dynamic pathway model of HGF-signal transduction, which allowed us to resolve the molecular mechanism resulting in reduced AKT phosphorylation in WD hepatocytes. A patient-adapted mathematical model correlated the basal MET phosphorylation with patient outcome after liver surgery and thus suggests MET phosphorylation as an indicator for liver disease burden.

## Results

### Characterization of proteomic alterations in Western Diet (WD) hepatocytes identifies dysregulated pathways

The development of non-alcoholic fatty liver disease (NAFLD) is characterized by the gradual accumulation of lipid droplets in hepatocytes. We hypothesized that these major structural changes have an impact on information processing and metabolic regulation in these cells. To induce a fatty liver-like phenotype, 8 weeks old C57BL/6N mice were fed with a high sugar high fat Western diet (WD) for up to 13 weeks. Age-matched mice fed with a standard diet (SD) served as controls. As expected, WD mice showed with a median weight of 39.4 g a significant increase in body weight, while SD mice remained at a median body weight of 29.7 g, (Fig 1A). Bright-field microscopy of primary mouse hepatocytes isolated from these mice (Fig 1B) revealed that lipid accumulation characterized by the formation of lipid droplets was indeed evident in the hepatocytes of the WD mice, but absent in those of the SD mice (Fig 1C, black arrows). To characterize the diet-induced changes in the protein composition of steatotic hepatocytes, we analyzed the proteome of both SD and WD hepatocytes by mass spectrometry (N=9 mice per condition) employing data independent acquisition (DIA). In total 4317 proteins were identified and a multidimensional scaling analysis provided evidence for major differences in the respective proteomes (Fig 1D). Data analysis utilizing *limma* (Ritchie *et al*, 2015), identified 301 proteins as differentially upregulated and 255 proteins as differentially downregulated in WD hepatocytes. To determine which pathways are primarily affected by feeding the WD, we performed an Ingenuity pathway analysis (Krämer *et al*, 2014) using significantly changed proteins as input (Benjamini and Hochberg adjusted p-value < 0.05 and log_2_ fold change < -0.5 or > 0.5). In Fig 1F, the enriched categories of canonical pathways with top 10 highest and lowest z- score are displayed. In line with previous studies (Greco *et al*, 2008; Puri *et al*., 2007) the majority of the identified pathways were connected to metabolism, including glucose, fatty acids and cholesterol metabolism. However, surprisingly the top most upregulated pathway was kinetochore metaphase signaling. The kinetochore is a complex of proteins responsible for anchoring the spindle fibers to duplicating chromatids and pulling sister chromatids during the metaphase of the cell cycle apart. Detected members of the kinetochore metaphase signaling pathways (Fig 1E) upregulated in WD hepatocytes were Macroh2a, H2ac20, H2az1 and H2ax, all linked to the nucleosome, and Zwint and Zw10, linked to kinetochore and mitosis, while Ppp1r10, a protein of the PTW/PP1 phosphatase complex controlling progression from mitosis to interphase, was downregulated. The HIF1α signaling pathway ranked second. The HIF1 α transcription factor has been linked in the context of the liver to lipid metabolism and vascular regulation during liver regeneration (Seo *et al*, 2020) and to enhanced proliferation of liver cancer cell lines (Tajima *et al*, 2009). Interestingly, the third most upregulated pathway in WD hepatocytes was ERK/MAPK signal transduction, a key pathway controlling proliferative responses. In sum, these proteomic alterations indicate that in steatotic hepatocytes the intricate network of metabolism and signal transduction controlling cell proliferation is dysregulated.

**Figure 1.**
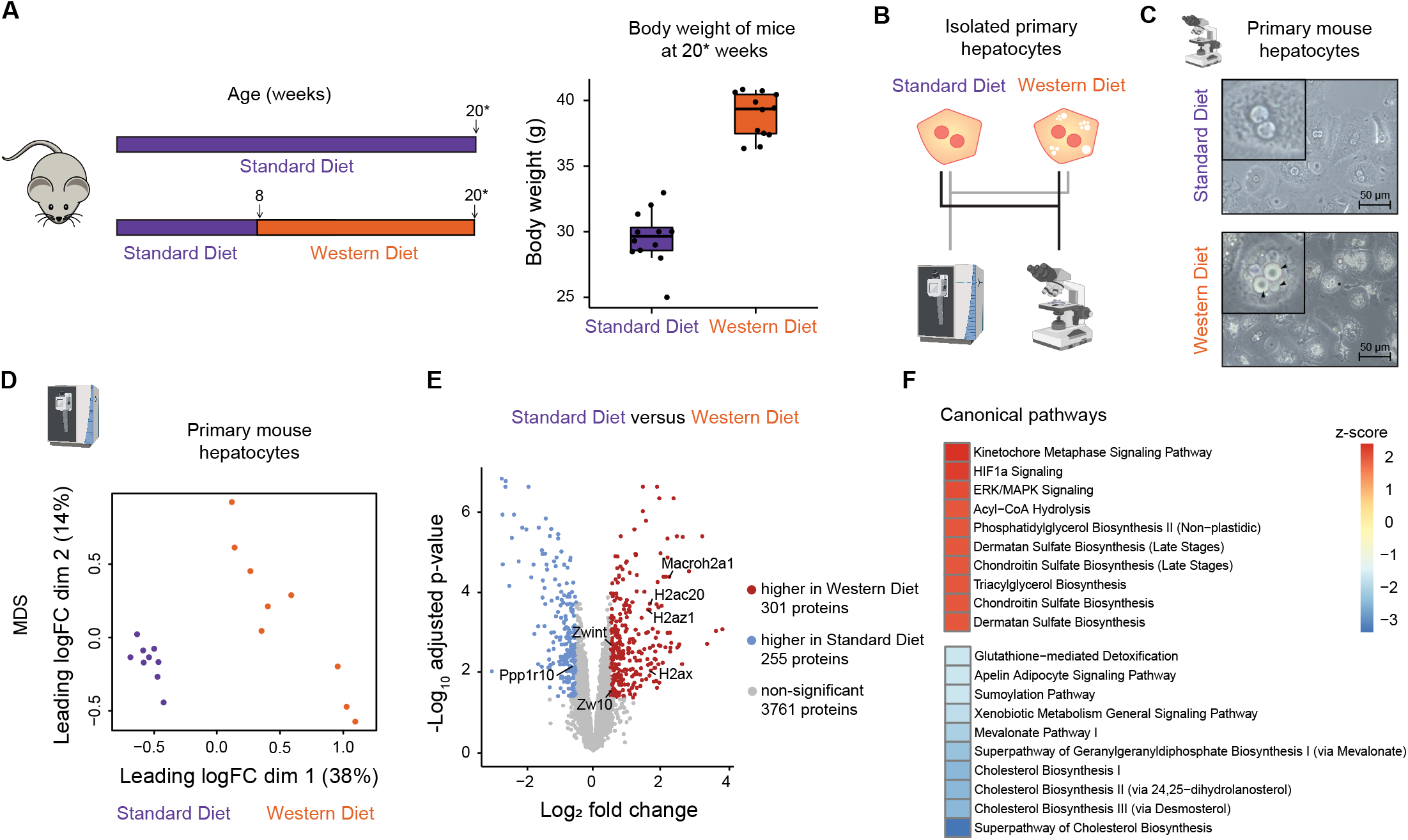
Western diet-induced proteome alterations. A. Schematic representation of the experimental setup. Mice were fed with standard diet (SD) for eight weeks and then switched to Western diet (WD) for 12 to 13 weeks or continued to receive SD as control. The body weight of SD and WD mice is shown as boxplot indicating median, 25th and 75th percentile. Dots represent data of single mice. 20*: mice have an age of 20 or 21 weeks. B. Primary mouse hepatocytes from SD and WD mice were isolated by liver perfusion, cultivated and characterized employing mass spectrometry and bright field imaging. C. Exemplary images from isolated primary hepatocytes depict lipid droplet formation in of hepatocytes derived from WD-fed mice. Arrows point to lipid droplets. D. Multidimensional scaling analysis of the hepatocyte proteome derived from SD and WD mice. Each dot represents the sample from one mouse. E. Up- and downregulated proteins were identified by log fold change and analysis of the adjusted p-value as depicted by the volcano plot. Proteins with fold change < -0.5 and p-value < 0.05, describing a downregulation in WD, are indicated in blue, proteins with fold change > 0.5 and adjusted p-value < 0.05, representing an upregulation in WD, are indicated in red. F. The top 10 up- and downregulated pathways in WD hepatocytes in comparison to SD hepatocytes are depicted as identified by Ingenuity pathway analysis (Krämer *et al*., 2014).

### Alterations in basal MET phosphorylation and AKT phosphorylation are characteristic features of WD hepatocytes

Hepatocytes in the liver are usually in a quiescent state and rarely proliferate in the absence of growth factors (Bottaro *et al*, 1991). The key growth factor controlling hepatocyte proliferation is HGF that binds to the receptor tyrosine kinase MET and triggers phosphorylation of Tyr 1232 and Tyr 1233 on the murine MET receptor (Tyr 1234 and Tyr 1235 on human MET). This enables the activation of PI3 kinase and MAP kinase signal transduction. To examine whether the uncovered proteomic alterations give rise to changes in the dynamics of HGF-signal transduction in WD hepatocytes, we designed based on our previous knowledge of the cross-talk of MAPK- and PI3K/AKT-pathways (D’Alessandro *et al*., 2015) and the link to proliferation (Mueller *et al*., 2015), dose- and time-resolved experiments to delineate by quantitative immunoblotting the differences in HGF-induced responses in WD and SD hepatocytes (Fig 2A). The HGF dose response experiments showed that in SD hepatocytes the HGF-induced phosphorylation of MET (double phosphorylation on Tyr 1232 and Tyr 1233) increased with higher doses and reached saturation at 40 ng/ml HGF (Fig 2B). Accordingly, the peak of HGF-induced phosphorylation of ERK (double phosphorylation on Thr 202 and Tyr 204 of ERK1/MAPK3 and on Thr 185 and Tyr 187 of ERK2/MAPK1) and of AKT (phosphorylation on Thr 308, referred as pAKT Thr 308 and phosphorylation in both Thr 308 and Ser 473, referred as ppAKT Ser 473 as the employed antibody detects exclusively the second phosphorylation of AKT) reached saturation at 40 ng/ml HGF. In WD hepatocytes at high HGF doses, a similar extent of MET and ERK phosphorylation was observed. In contrast, the phosphorylation at basal level and the phosphorylation of MET upon low doses of HGF were higher in WD hepatocytes when compared to SD hepatocytes. Even more strikingly, at all HGF doses higher than 1 ng/ml, AKT phosphorylation was significantly reduced in WD hepatocytes. The comparison of the dynamics of HGF-induced signal transduction in SD and WD hepatocytes (Fig 2C) revealed that the peak amplitude of the tyrosine kinase receptor MET was higher in SD hepatocytes, while the basal amount of phosphorylated MET was elevated in WD hepatocytes. In line with the notion that ligand mediated activation of MET induces its internalization and degradation (Jeffers *et al*, 1997), we observed a decline of total MET after HGF stimulation in SD hepatocytes, but this decline was reduced in WD hepatocytes. As expected, the phosphorylation of ERK correlated in SD and WD hepatocytes with the respective phosphorylation dynamics of MET. Interestingly, although the total amount of AKT was comparable between SD and WD hepatocytes, in line with the results obtained in the dose response experiment (Fig. 2B), HGF-induced phosphorylation of AKT was much reduced in WD hepatocytes compared to those from SD mice (Fig. 2C). In conclusion, our results suggested that exposure of hepatocytes to a high sugar and high fat diet results in a dysregulation of HGF-induced proliferative signal transduction.

**Figure 2.**
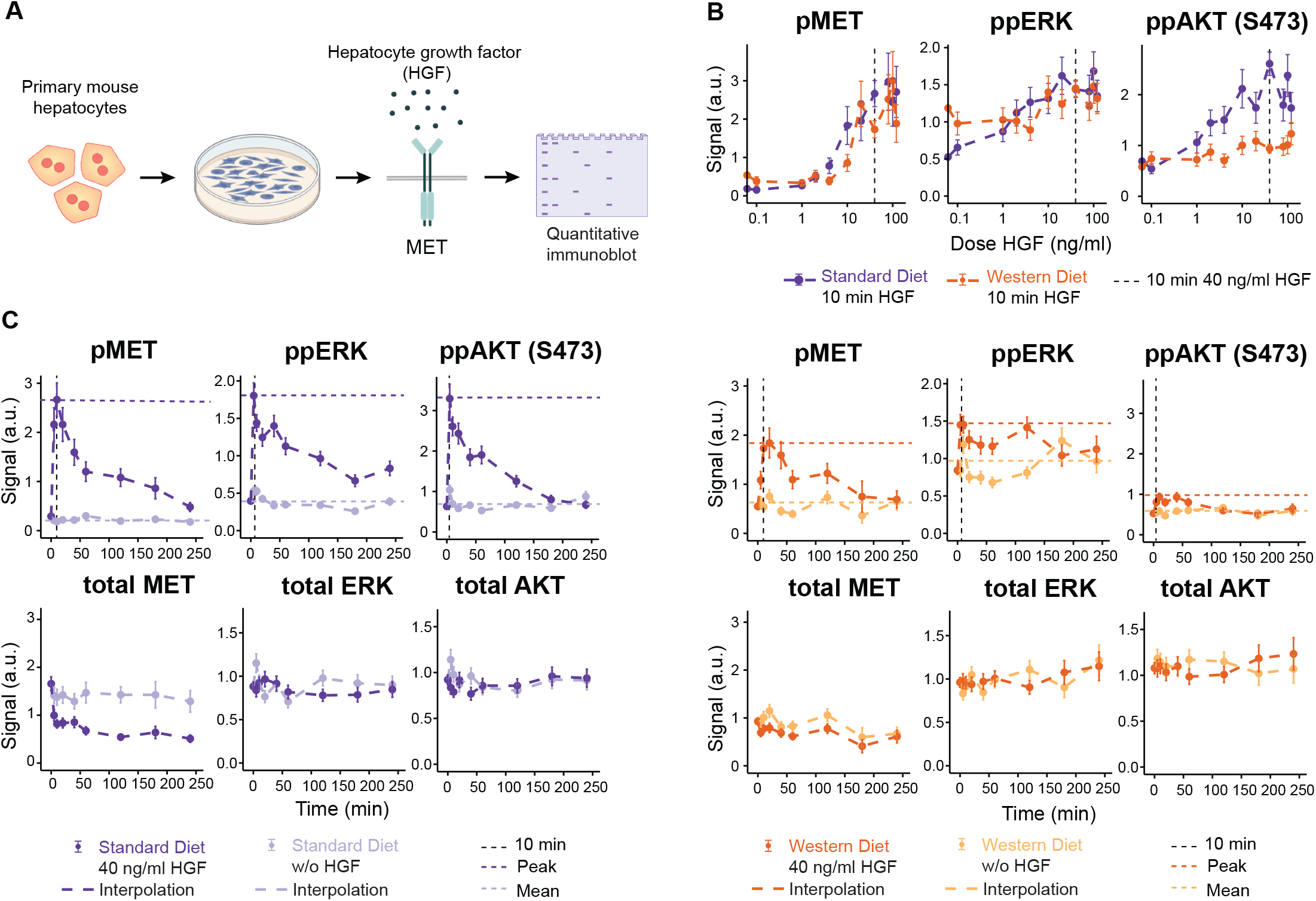
HGF-induced activation of signal transduction in SD and WD hepatocytes. A. Experimental design of HGF stimulation experiments in primary mouse hepatocytes. Cells were seeded 24h prior to the start of the experiment. 18 to 20h before stimulation, cells were transferred to a serum-free medium and switched to serum- and dexametasone-free medium 6 hours prior to the stimulation with 40 ng/ml HGF, or were left untreated. Measurements were taken at indicated time points. B. HGF dose dependency of MET, ERK and AKT phosphorylation in primary SD and WD mouse hepatocytes. Cells were stimulated with indicated doses of HGF for ten minutes and phosphorylation of MET and ERK and AKT was quantified by immunoblotting. Data points are displayed as dots with 1*σ* confidence interval estimated from biological replicates (N = 3-9) using a combined scaling and error model. Solid lines represent linear interpolations. C. Time course measurements of HGF-induced signal transduction in primary mouse hepatocytes of SD and WD mice. Cells were stimulated with 40 ng/ml HGF for up to 4 hours and the phosphorylation as well as the abundance of MET, ERK and AKT was quantified by immunoblotting. Data points are displayed as dots with 1*σ* confidence interval estimated from biological replicates (N = 3 to 9) using a combined scaling and error model. Dashed lines indicate basal and peak signal levels.

### Dynamic pathway model-based analysis identifies basal MET phosphorylation and protein abundances as dysregulated in primary mouse hepatocytes

To unravel the molecular mechanism leading to decreased HGF-induced AKT phosphorylation in WD hepatocytes, we developed a mechanistic mathematical model of HGF-induced signal transduction including the nutrient sensor mTOR that modulates MAPK and PI3K/AKT signal transduction as a function of the available metabolites. We hypothesized that the high sugar and high fat diet affects mTOR signal transduction and as a consequence impacts the activation of the pro-mitogenic ribosomal protein S6 as well as the phosphorylation of ATK (Fig 3A).

**Figure 3.**
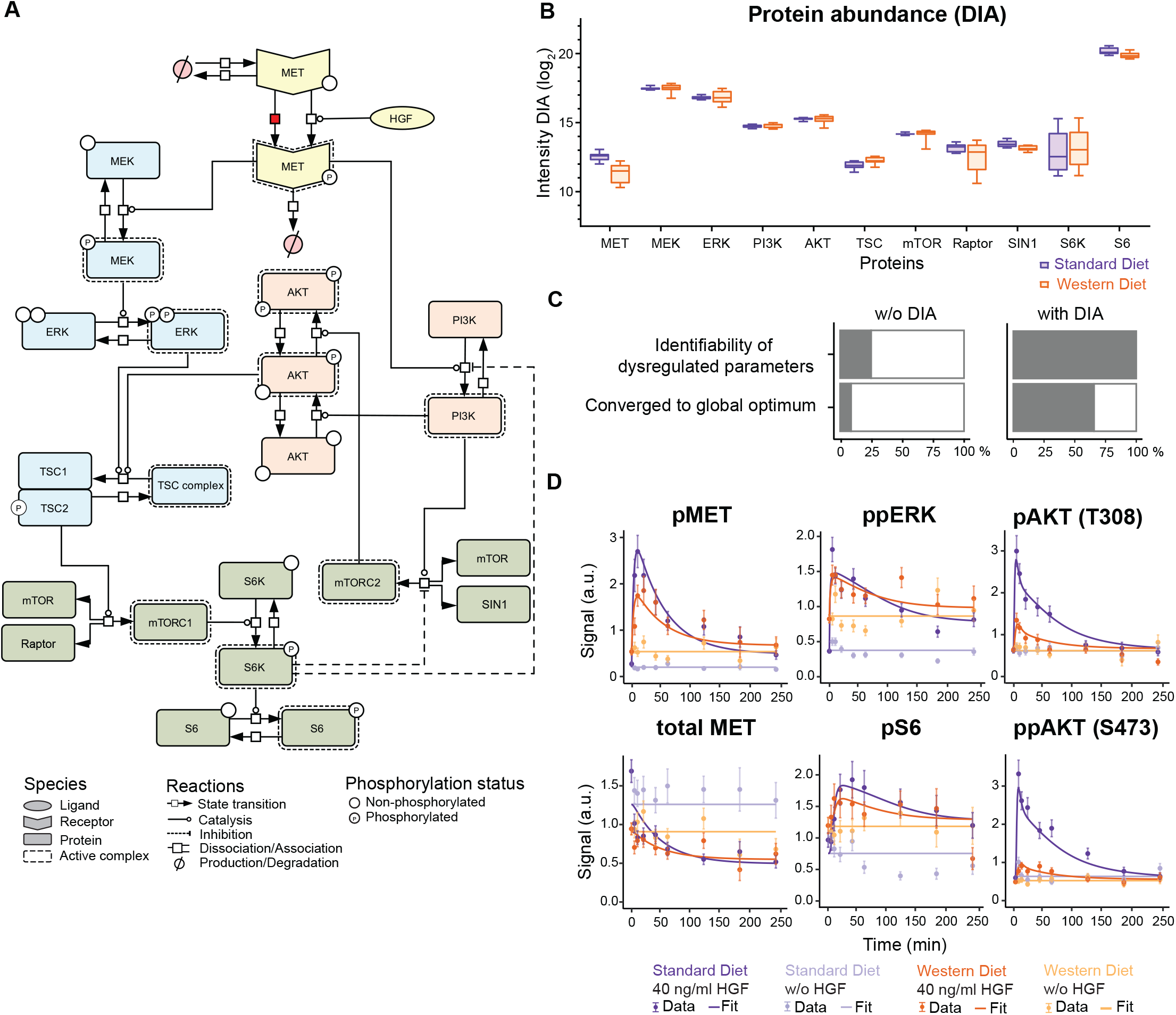
Modeling WD-induced alterations in HGF signal transduction. A. The structure of the mathematical model capturing HGF-induced signal transduction via the MAPK cascade (blue), the PI3K pathway (red) and mTOR signaling (green) is displayed according to Systems Biology Graphical Notation (Le Novere *et al*, 2009). All parameter values were implemented as identical for WD and SD hepatocytes except for the basal MET phosphorylation rate, indicated by the red box, and protein abundances. B. Measurements of protein abundances derived from primary mouse hepatocytes. Lysates of unstimulated hepatocytes were subjected to data-independent mass spectrometry analysis. Resulting data was LFQ normalized and represented as boxplot. C. Impact of the DIA data on parameter identifiability and convergence. The identifiability of the twelve dysregulated parameters increases from 25% to 100% upon DIA incorporation, while the convergence to the global optimum during optimization increases from 9% to 66%. D. Model calibration with time-resolved immunoblot measurements for MET, ERK, S6 and AKT phosphorylation and MET abundance upon stimulation with 40 ng/ml HGF. Data points are displayed as dots along with 1*σ* confidence interval estimated from biological replicates (N = 3 to 9) using a combined scaling and error model. Model trajectories are depicted as solid lines.

The dynamic pathway model is based on coupled ordinary differential equations (ODEs) and describes the interconnection between HGF-induced signal transduction and mTOR. In the mathematical model, MET is subject to constant production and degradation, while phosphorylated MET is degraded with a different rate. The signal emanating from the activated MET receptor is propagated by two distinct signal transduction pathways, MAPK and PI3K/AKT. In the MAPK cascade, phosphorylated MET leads to MEK phosphorylation, which in turn phosphorylates ERK. In the PI3K pathway, phosphorylated MET recruits PI3K, which induces the first phosphorylation of AKT, at Thr 308 (pAKT). Both phosphorylated ERK and AKT phosphorylated on Thr 308 can inactivate the TSC complex by phosphorylating TSC2, which allows mTORC1 formation and activation. Active mTORC1 phosphorylates S6K that in turn activates the pro-mitogenic ribosomal protein S6. Additionally, activated PI3K induces mTORC2 complex formation, which leads to phosphorylation of AKT on the second phosphorylation site Ser 473 resulting in AKT phosphorylated on Thr 308 and Ser 473 (ppAKT) and serving as a readout for the activation of the mTORC2 complex. Two negative feedback loops originating from phosphorylated S6K are represented in the mathematical model in a condensed manner (Fig 3A, dashed lines). First, activated S6K leads to the inhibition of the mTORC2 complex. Second, activated S6K phosphorylates and inhibits IRS1, which prevents activation of PI3K. In sum, our mathematical model includes 26 ordinary differential equations (for model reactions and observation functions, see Dataset EV1 and Dataset EV2) and 23 different model states.

To capture the dynamic properties of the system, the parameters of the mathematical model were calibrated based on the quantitative dose- and time-resolved immunoblot data shown in Fig 2. Additionally, we generated data on the HGF dose-dependent phosphorylation of AKT on the phosphorylation site Thr 308 (pAKT) as well as on the amount of total MET, total AKT and total ERK (Fig EV1A, data points). Further, the time-resolved dynamics of AKT phosphorylation on Thr 308 and of pS6 (Fig 3D, data points) and of total S6 (Fig EV1B, data points) were recorded. In total, 491 data points generated under 22 experimental conditions were employed for the calibration of our mathematical model. Since previous model-based investigations revealed that differences in the basal protein abundance greatly affect the dynamics of information processing (Adlung *et al*, 2017), we assumed as a start that all basal protein levels were different in SD and WD hepatocytes, and thus were considered as dysregulated parameters. To investigate if, in addition, dynamic parameters such as phosphorylation rates were affected by the Western diet, we performed a comparison of different model hypotheses by evaluating the resulting model fits based on the Bayesian information criterion (BIC) (Fig EV1C). Interestingly, our analysis revealed that all but one dynamic parameter could be assumed as identical retaining a good fit to the data of SD and WD hepatocytes. For the accurate model-based representation of the data for both conditions it is necessary and sufficient if only the dynamic parameter for the basal (HGF-independent) phosphorylation rate of MET is increased in WD hepatocytes (Fig 3A, reaction marked in red).

Thus, twelve dysregulated parameters appear to be required to describe the altered HGF-induced signal transduction in primary mouse hepatocytes induced by the Western diet, comprising the basal phosphorylation rate of MET and differences in the abundance of eleven proteins. To assess the identifiability of these twelve dysregulated parameters, we calculated profile likelihood-based confidence intervals (Raue *et al*, 2009) for each parameter (Fig EV2A). This analysis showed that only three of these parameters where identifiable for both conditions, i.e. had defined confidence intervals (Fig. 3C). Therefore, additional experimental data was required to better specify the protein abundance in both SD and WD hepatocytes. Accordingly, we examined total protein lysates of SD and WD hepatocytes by global proteomics employing mass spectrometric analysis operated in the data independent acquisition (DIA) mode. We detected a significant decrease in the abundance of MET and a decrease of S6 in the WD hepatocytes, while the abundance of the other protein species was comparable between SD and WD hepatocytes (Fig 3B). Because global proteomics only provides relative intensities that approximately scale with the concentration of the measured proteins, we reasoned that we had to acquire quantitative values on the abundance of at least one protein species and implement this data set as a reference in the mathematical model. Therefore, we determined the abundance of AKT using quantitative immunoblotting (Fig EV3A). This measurement and the knowledge of the average protein content of hepatocytes allowed us to estimate the number of AKT molecules per cell, which was included as additional information in the mathematical model. By linking the relative amount of AKT protein determined by our mass spectrometry-based DIA measurements with the corresponding absolute amount of AKT molecules per cell, the model could infer the corresponding protein concentrations for all proteins of interest. With this additional information it was possible to estimate all twelve dysregulated parameters with narrow confidence intervals in both conditions (Fig EV2B). Importantly, the inclusion of the absolute values for the protein abundance of SD and WD hepatocytes increased the identifiability of the dysregulated parameters from 25% to 100% and the convergence to the global optimum from 9% to 66% (Fig 3C, Fig EV2C). As a result, the final mathematical model, which included only one diet-specific dynamic parameter, the basal MET phosphorylation rate, and 11 diet-specific parameters for protein abundance, was able to explain the increased basal phosphorylation and reduced amplitude of pMET and pERK as well as the reduced phosphorylation of AKT in response to HGF stimulation in WD hepatocytes (Fig 3D). Further, the model trajectories were in agreement with the experimentally observed HGF dose-dependent dynamics of pMET, pERK, pAKT (Thr 308) and ppAKT (Thr 308, Ser 473) in WD and SD hepatocytes (Fig EV1A). The final mathematical model was capable of capturing the dynamics of total MET (Fig 3D) as well as the differences in the total protein amounts of the analyzed proteins (Fig EV1A-B) and was in line with the protein concentrations determined by mass spectrometry-based DIA measurements in SD and WD hepatocytes (Fig EV3B).

In sum, we established a mathematical model of HGF-signal transduction that was able to explain the alterations in HGF-induced MET and AKT phosphorylation dynamics in WD hepatocytes (for estimated parameter values, see Dataset EV3 and Dataset EV4). Importantly, the inclusion of the metabolite sensing pathway mTOR, in form of a negative feedback loop between mTORC1 and mTORC2, proved to be essential to gain insights into the molecular mechanisms causing the reduced AKT phosphorylation in WD hepatocytes: The increased basal MET phosphorylation in WD hepatocytes results in an increased basal phosphorylation of S6K, which in turn inhibits mTORC2 and PI3K activation and as a consequence, results in reduced AKT phosphorylation.

### The basal phosphorylation rate parameter of MET is sufficient to explain altered HGF- signal transduction in WD hepatocytes

To investigate which of the twelve diet-specific parameters had the largest impact on the observed WD-specific dynamics of HGF induced signal transduction, we performed a parameter scan. By fixing all but one of the dysregulated parameters to the values estimated for SD hepatocytes, we gradually shifted the value of the dysregulated parameter of interest from the SD estimated value to the WD estimated value. As the main differences in HGF- signal transduction in WD hepatocytes compared to SD hepatocytes, we identified increased basal phosphorylation of MET and ERK and decreased HGF-induced phosphorylation of AKT on Thr 308 and Ser 473. Therefore, we selected these as metrics to assess the impact of each parameter on HGF-signal transduction and simulated the trajectories of pMET, pERK and ppAKT (Thr 308, Ser 473) for the different parameter values. The model simulations (Fig 4A and Fig EV4) showed the largest effects when altering the basal protein abundance of MET *(k_total MET_),* MEK *(k_total MEK_)* and S6K *(k_total S6K_)* as well as the basal phosphorylation rate of MET *(k_basal p-rate MET_).* Solely by shifting the basal MET phosphorylation rate to the value determined for WD hepatocytes it was possible to reproduce the increased basal phosphorylation of MET and ERK, as well as the decreased AKT activity leading to reduced area under the curve (AUC) of ppAKT (Thr 308, Ser 473) as previously observed in WD hepatocytes. To quantitatively asses these three features, we re-optimized each dysregulated parameter individually, in the range between SD and WD estimates, to best match the amount of basal pMET, basal ppERK and the AUC of ppAKT (Thr 308, Ser 473) as observed in WD hepatocytes. In line with the results obtained by the parameter scan (Fig. 4A), the only parameter that enabled the reproduction of the WD-specific features was the basal MET phosphorylation rate (Fig 4B), supporting its key role as driver for the WD-specific alterations in HGF-induced signal transduction.

**Figure 4.**
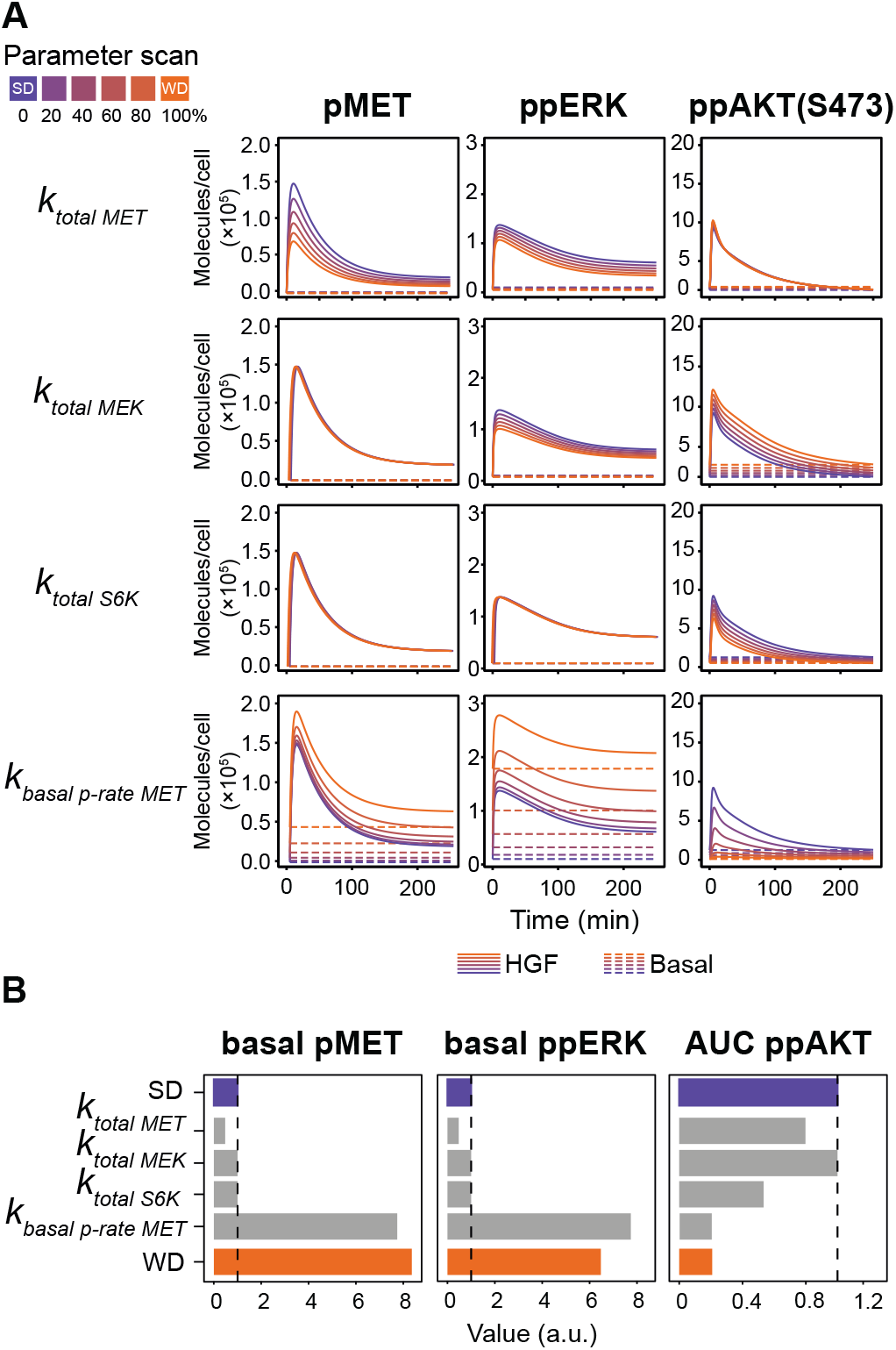
Influence of dysregulated parameters on protein dynamics. A. Phosphorylation dynamics of MET, ERK and AKT are displayed as simulated based on a parameter set with parameter values fixed to the SD estimates for all but one of the dysregulated parameters. The value for the indicated dysregulated parameter was gradually shifted from the SD estimate (purple) to the WD estimate (orange), simulating protein dynamics for each step. The influence of those dysregulated parameters with the biggest effect on model trajectories are shown. Solid lines indicate model trajectories after HGF stimulation and dashed lines indicate the control condition. B. Quantitative analysis of the ability of dysregulated parameters to reproduce WD-specific features. We reoptimized each dysregulated parameter individually in the range between SD and WD estimates to determine the best fit for the three features. Colored bars represent the feature value as determined from the original model fit for SD and WD. Grey bars indicate the optimized feature value for dysregulated parameters.

### WD hepatocytes show an increased proliferative behavior

As a consequence of the increased basal MET phosphorylation rate, we observed an elevated basal phosphorylation of the pro-mitogenic ribosomal protein S6 in WD hepatocytes. Therefore, we hypothesized that these changes might influence proliferative responses in the presence and absence of HGF. To explore this hypothesis, we fed 8 week old mice expressing the Fluorescent Ubiquitination-based Cell Cycle Indicator (Fucci2) (Abe *et al*, 2013) with either SD or WD for 12 weeks and isolated primary murine hepatocytes (Fig 5A). For each condition, we monitored ten individual hepatocytes derived from three SD and three WD mice, respectively, by live cell microscopy and acquired time-resolved data on the changes in fluorescence intensities of the FUCCI system for up to 65 hours (Fig 5B). To quantify the cell-cycle entries as a measure for proliferative behavior, we performed single-cell tracking and counted cell cycle entries of each hepatocyte defined as the transition from S-phase to G2/M phase (Fig 5C). For 13 out of 30 WD hepatocytes (43%) cell cycle entry was observed within the observation time, whereas only 7 out of 30 SD hepatocytes (23%) showed such a response. This observation indicates that already in the absence of HGF there is a marked increase in cell cycle entries in WD compared to SD hepatocytes. Upon HGF stimulation, all tracked SD and WD hepatocytes showed cell cycle progression, confirming that growth factor responsiveness was maintained in WD hepatocytes. Importantly, in response to HGF stimulation 28 out of 30 WD hepatocytes (94%) underwent two to four rounds of cell cycle progression within the observation time. In contrast, only 11 out of 30 SD hepatocytes (36%) showed two or three rounds of cell cycle progression. Taken together, these results revealed that proliferative responses are enhanced in WD hepatocytes even in the absence of HGF. These observations imply that the identified increase in basal MET phosphorylation in WD hepatocytes is an indicator of altered signal transduction enabling HGF-independent proliferation.

**Figure 5.**
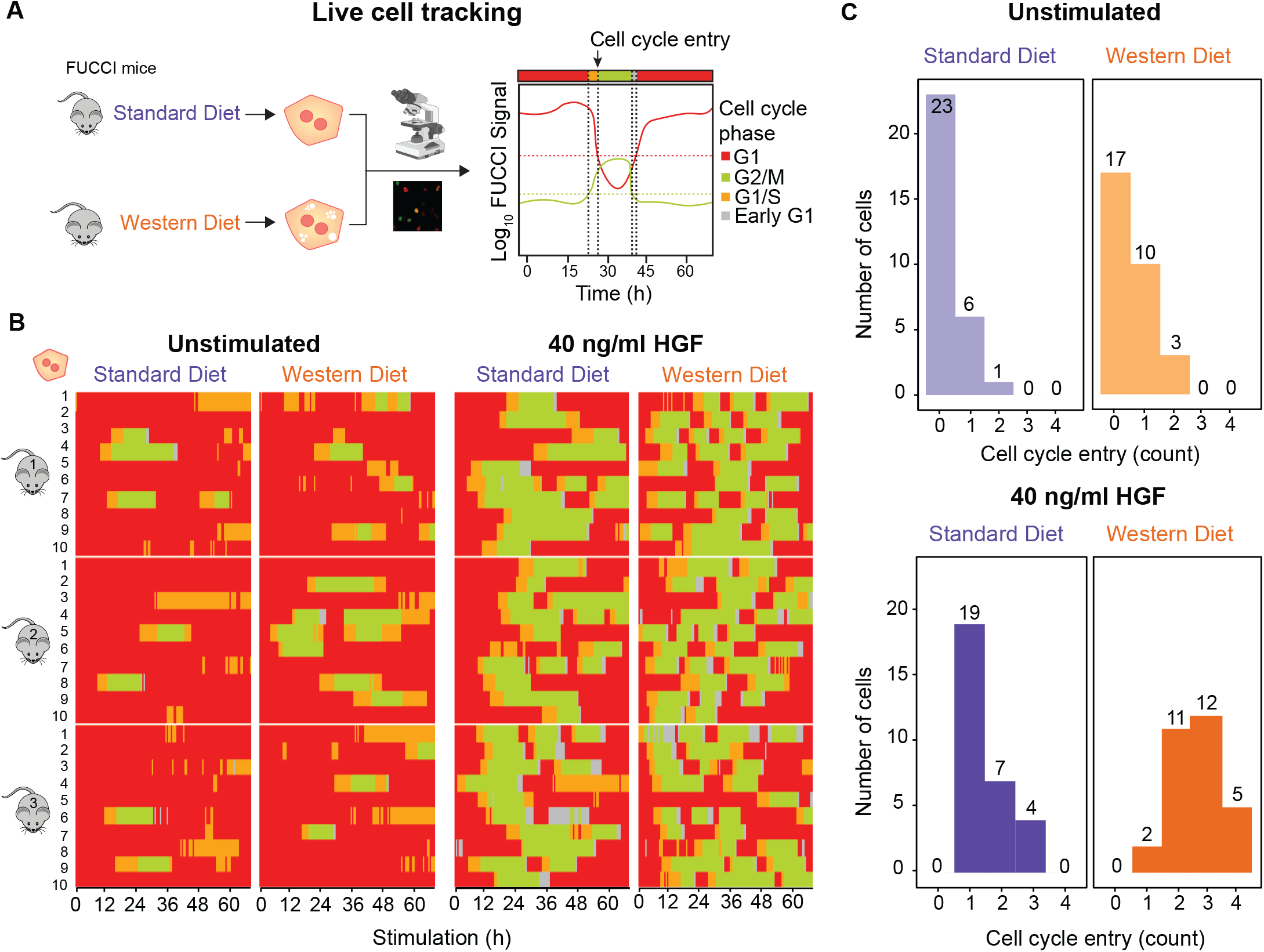
Altered proliferation of WD hepatocytes. A. SD and WD mice carrying the Fucci2 cell cycle reporter were used to track cell cycle entries of primary hepatocytes via live cell imaging. Cells were transduced with adeno-associated viral vectors encoding Histone2B–mCerulean to enable tracking of the cells and assigned to be in a given cell cycle phase based on threshold defined for the FUCCI signals. B. Live cell microscopy was performed with sampling rate of 15 min for up to 65 h, and ten exemplary hepatocytes of three SD and three WD mice were either stimulated with 40 ng/ml HGF or left untreated. The time-dependent cell cycle phases G1, G1/S, and S/G2/M and early G1 are displayed for each cell. C. Quantification of cell cycle entries per cell calculated from the single cell analysis shown in (B). A cell cycle entry was considered if cells transited from S to G2-phase.

### Basal MET phosphorylation is indicative for liver disease burden in patient-derived hepatocytes

Our observation that basal MET acts as an integrator of structural and metabolic alterations during the progression of chronic liver disease in mice, let us to propose that basal MET phosphorylation could be a useful indicator for the burden of liver disease in humans. To test this hypothesis, we analyzed HGF-induced signal transduction in patient-derived primary human hepatocytes isolated from tumor-free tissue of seven patients with different liver pathologies that underwent partial liver hepatectomy (see Dataset EV5 for patient anamnesis). The primary human hepatocytes were stimulated with HGF and cellular lysates taken at time points up to 120 min were analyzed by quantitative immunoblotting to generate time-resolved information on the dynamics of HGF-induced signal transduction in the hepatocytes of the individual patients (Fig. 6A) In addition, the proliferation behavior was also determined (Fig EV5A). We examined the HGF-induced phosphorylation dynamics of MET (Tyr 1234 and Tyr 1235), ERK (Thr 202 and Tyr 204 on ERK1, Thr 185 and Tyr 187 on ERK2), AKT (Ser 473) (Fig 6A, data points) and S6K (Thr 389) in the patient-derived hepatocytes (Fig EV5B). Likewise, the total amounts of MET, ERK, AKT and S6K were determined by quantitative immunoblotting (Fig EV5B) and complemented by protein abundances quantified by DIA-based mass spectrometry (Fig EV5C, box plots). Our dynamic pathway model for HGF-induced signal transduction developed for murine SD and WD-derived hepatocytes was adapted to analyze the time-resolved data obtained for the patient-derived hepatocytes. The majority of the model parameters was kept as estimated for the murine system. The model parameters identified as dysregulated in the hepatocytes from WD mice, comprising protein abundances and the basal MET phosphorylation rate, were adapted to the conditions in the hepatocytes of individual patients. The model parameters identified as dysregulated in the hepatocytes from WD mice, comprising protein abundances and the basal MET phosphorylation rate, were adapted to the conditions in the hepatocytes of individual patients. However, this model was yet insufficient to adequately describe the human data (98 parameters, BIC = 805). Therefore, we included in addition as human-specific parameters the HGF-induced MET phosphorylation rate as well as the degradation rate of the activated MET receptor (100 parameters, BIC = 564). These findings are supported by previous observations reporting a difference in the binding affinity of HGF to MET between mice and humans (Bussolino *et al*, 1992), which affects HGF binding to the receptor and as a consequence receptor phosphorylation and degradation of the ligand-receptor complex. In total 852 data points were used for the calibration of 100 human- and patient-specific parameters (see Dataset EV6 for estimated model parameters of primary human hepatocytes). The calibrated dynamic pathway model of HGF signal transduction in primary human hepatocytes was able to capture the basal (Fig 6A and EV5B, dashed lines) as well as the HGF-induced patient-specific dynamics (Fig 6A and EV5B, solid lines) of all pathway components. Further, the model fits for total protein abundances were in line with the global proteome measurements acquired by DIA-mass spectrometry (Fig EV5C, points). Both experimental data and model trajectories revealed a patient-to-patient difference in the basal MET phosphorylation levels of untreated primary human hepatocytes (Fig 6A). Since our model-based analyses of HGF signal transduction in hepatocytes of mice with WD-induced chronic liver disease suggested a relation between the basal MET phosphorylation level and disease burden, we investigated the information encoded in the dynamics of HGF-induced signal transduction in patients. To this aim, we utilized our mathematical model to calculate the patient-specific basal MET phosphorylation rate, the AUC of ppAKT and the MET abundance. In addition, we introduced ratios of these characteristic features accounting for interconnected complexity. We also included the quantification of the HGF-induced proliferation of the patient-derived human hepatocytes as a proxy for the proliferation potential at the organ level. To test how informative the model-based characteristics and these measurements are, we correlated them to several patient-specific clinical features (Dataset EV7), which we divided in three subgroups: (i) Pre- and post-operative blood metrics: hepatocyte growth factor (HGF), Interleukin 6 (IL6), Interleukin 8 (IL8), Platelet-derived growth factor (PDGF), Tumor growth factor beta 1 (TGFβ1) and Tumor necrosis factor alpha (TNFα); (ii) patient features, such as the age, the body mass index (BMI), the Fibrosis score and the Charlson Comorbidity Index (CCI); and (iii) the patient outcome including intensive care and hospitalization days, the Clavien Dindo score and the complication index. Surprisingly, the *ex vivo* proliferation of hepatocytes did not correlate with the patient outcome, emphasizing the complexity of the proliferative response in the liver (Fig 6B). The AUC of ppAKT as well as the corresponding ratio *k_basal p-rate MET_* /*AUC _ppAKT_* did not correlate with the patient outcome but to the post-operative concentrations of HGF, IL8 and TNFα. However, the basal MET phosphorylation rate, *k_basal p-rate MET_*, showed a significant correlation with three of four patient outcome measures. This correlation was also present for the ratio *k_basal p-rate MET_* /*k _total MET_*, but to a smaller extend. Of the six factors, HGF, IL6, IL8, PDGF, TGFβ1 and TNFα longitudinally determined in the blood, only PDGF showed correlation with *k_basal p-rate MET_* at all time points. Taken together our results suggested that *k_basal p-rate MET_* and *k_basal p-rate MET_* /*k _total MET_* might provide as suitable measure to predict patient outcome.

**Figure 6.**
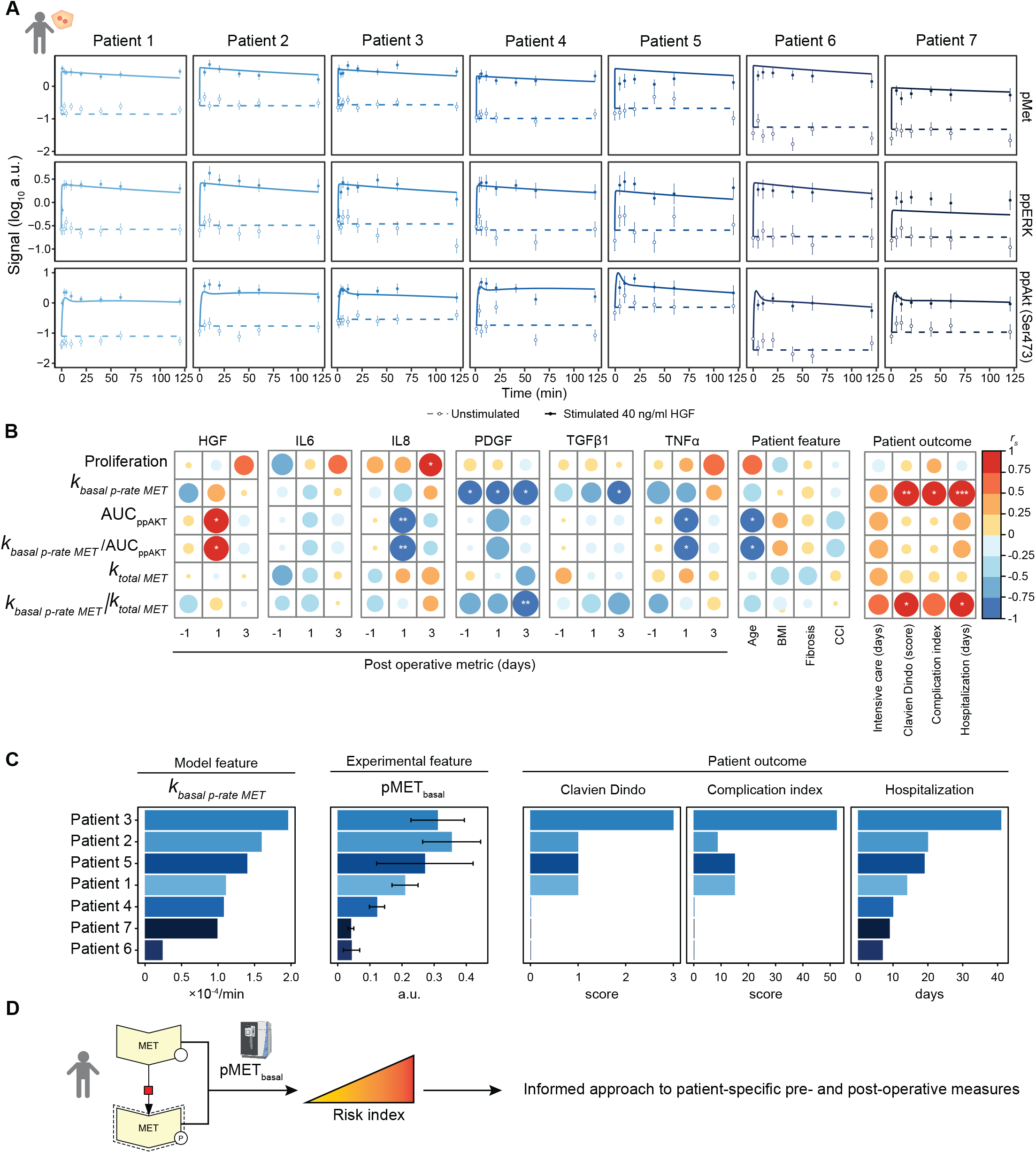
Basal pMET levels in primary human hepatocytes correlate to patient outcome. A. Time-resolved immunoblot measurements and model fits for pMET, pERK and ppAKT in primary human hepatocytes derived from seven patients and stimulated with 40 ng/ml HGF. Data points are displayed as dots along with 1*σ* confidence interval estimated from biological replicates (N = 1 to 3 per patient) using a combined scaling and error model. Model trajectories are depicted as lines. B. Spearman correlation of hepatocyte proliferation and model features with patient-specific characteristics, blood metrics and outcome. Significance levels are indicated as * p < 0.05, ** p < 0.01, *** p < 0.001. BMI: body mass index, CCI: Charlson comorbidity index. C. The patient-specific basal MET phosphorylation rate as estimated by the model is depicted in comparison to the measured basal pMET levels, obtained as mean of all unstimulated pMET measurements per patient. In comparison, the clinical metrics Clavien Dindo, complication index and hospitalization are shown. Patients are sorted by the basal MET phosphorylation rate, colors indicate patient number. Error bars represent the standard deviation of 7 to 9 replicates. D. Proposed use of the basal phosphorylation of MET as an informative metric for liver disease burden and patient recovery after liver surgery.

As the parameter estimation of *k_basal p-rate MET_* cannot be easily introduced in a clinical setup, we tested if the basal phosphorylation of MET, which was used to parametrize the mathematical model, could describe the patient outcome. In Fig 6C, the patients were sorted by the model-derived estimates of the basal MET phosphorylation rate regarding potential clinical outcome. The values of pMET that were experimentally determined by quantitative immunoblotting for each patient, corroborated with the order of the patients based on the basal MET phosphorylation rate. In line with the correlation analysis, Clavien Dindo score, complication index and especially hospitalization time directly correlated with both model-derived basal MET phosphorylation rate and experimentally measured basal pMET levels. In conclusion, our results identified the basal phosphorylation of MET as key node integrating alterations in hepatocytes and suggest it as an informative metric (Fig 6D) for liver disease burden and patient recovery after liver surgery.

## Discussion

In this study, we characterize Western diet-induced phenotypic changes in murine hepatocytes and employ dynamic pathway modelling to resolve the underlying molecular mechanisms resulting in altered intracellular signal transduction and enhanced proliferation of WD hepatocytes. A key observation was that the basal MET phosphorylation is increased in WD mice after 12 weeks of WD-feeding, while conversely the amplitude of the MET phosphorylation dynamic is reduced. These insights required the generation of high-quality quantitative data on the dynamics of HGF signal transduction and thus shed new light on the complex regulation of MET in NAFLD, while MET was so far primarily studied in the context of liver regeneration or cancer (Bottaro *et al*., 1991; Paranjpe *et al*., 2016). With our mathematical modelling approach, we resolve that the elevated levels of basal MET phosphorylation result in surprisingly strong inhibition of AKT phosphorylation in WD hepatocytes. Conversely, it was reported in the context of type II diabetes and fatty liver disease that increased phosphorylation of AKT was an indicator of insulin response and consequently a good prognostic marker for reduced steatosis (Vivero *et al*, 2021). Of note, the known interactions between MET and the insulin receptor substrate 1 (IRS1) (DeAngelis *et al*, 2010) have not been explicitly included in our model, but could be integrated in the future to further elucidate effects resulting in the observed downregulation of AKT as IRS1 is a target protein of the negative feedback regulation via S6K (Tremblay *et al*, 2007; Zhang *et al*, 2008).

The development of mechanistic models provides a tool to understand complex biological questions and to gain insights into molecular mechanisms regulating dynamic behavior. Accordingly, based on mathematical modeling of the HGF signal transduction pathway cross-talk, we dissected dependencies of MAP kinase and PI3 kinase signal transduction in primary hepatocytes and established the link to the regulation of cell cycle progression (D’Alessandro *et al*., 2015; Mueller *et al*., 2015). Further, Dalla Pezze et al. resolved the interactions of the mTOR pathway upon insulin stimulation (Dalle Pezze *et al*, 2012). Based on these findings, we included in our model mTOR as a metabolic gateway in the liver (Jia *et al*., 2014) to capture the Western diet-induced alterations in HGF-dependent signal transduction in primary hepatocytes. Interestingly, the only reaction rate, which we identified with our mathematical modelling approach as dysregulated in this disease scenario, was the basal phosphorylation of the MET receptor. Further analysis of the mathematical model showed that the reaction rate of basal MET phosphorylation was sufficient to describe the observed diet-induced changes in AKT phosphorylation and MAPK signal transduction. These results disentangle the complex interrelations of the impact of metabolic alterations and proliferative signal transduction in hepatocytes and thus shed new light on molecular dysregulation in the context of chronic liver diseases. So far, the analysis of HGF-induced signal transduction in the liver has primarily focused on its role in repair of liver damage and regeneration (Oe *et al*., 2005), whereas the impact of metabolic alterations, which could be critical at early stages of chronic liver diseases, has not yet been considered. Importantly, we uncover that the exposure to the Western diet increases not only the levels of basal MET phosphorylation but also results in an increased proliferation of hepatocytes. These insights suggest that although the early steatotic phenotype in liver disease may be reversible, the metabolic state of the hepatocytes and their capacity to interpret external signals has shifted. We have not directly addressed the molecular mechanism driving the increased basal phosphorylation of MET, but hypothesize that the diet-induced changes in the bilipid layer of the hepatocytes may alter the affinity and the dimerization rate of MET rendering the receptor a central integrator of metabolic changes. In support of this hypothesis, it has been reported that a high fat diet decreased the cholesterol content of the plasma membrane of hepatocytes and reduced the affinity of the insulin receptor to insulin (Sabapathy *et al*, 2022). In line with this report, our analysis of canonical pathways showed that cholesterol biosynthesis was downregulated in WD-derived hepatocytes. Since HGF has been identified as modulator of liver fibrosis, which develops as sustained liver disease (Kwiecinski *et al*, 2011; Tekkesin *et al*., 2011), we propose that HGF-induced signal transduction determines the fate of hepatocytes and based on our results the extent of basal MET phosphorylation could be a central indicator facilitating the quantification of the alterations.

Previous reports indicated that liver steatosis affects the regenerative capacity of the liver (Allaire & Gilgenkrantz, 2018; Ghanemi *et al*, 2020). Therefore, our observations could be of relevance for patients undergoing liver surgery. To assess this, we analyzed primary hepatocytes isolated from seven patients undergoing liver hepatectomy to adapt our dynamic pathway model of HGF signal transduction to the human situation. In line with our previous studies (Dehlke *et al*, 2022; Murtha-Lemekhova *et al*, 2021), we observed little correlation (p>0.01) of HGF, IL6, IL8, PDGF, TGFβ1, TNFα, age, BMI, and fibrosis with the patient outcome, which was assessed by Clavien Dindo score, complication index and hospitalization days. In contrast, we uncovered that the basal phosphorylation rate of MET strongly correlated with the patient outcome. The importance of the basal MET phosphorylation rate in this context was further supported by the immunoblot based quantifications of the pMET level in the patient-derived hepatocytes that followed the trend of the basal MET phosphorylation rate and the clinical outcome. Furthermore, it has been reported that hepatocytes undergo metabolic reprograming during liver proliferation (Chembazhi *et al*, 2021). It was proposed that since proliferating hepatocytes cannot sustain liver specific metabolic functions, other hepatocytes shift into a hyperactive metabolic state. These notions could explain why patients, with higher basal MET phosphorylation, undergoing hepatectomy are less capable to compensate the metabolic function in the liver and show higher risk of hepatic failure. Nevertheless, we acknowledge that a cohort of seven patients is very limited and our observations should be interpreted with caution.

In conclusion, our model-based insights identify the basal phosphorylation rate of MET as an indicator of alterations in the metabolic state of hepatocytes and its impact on proliferative signal transduction. Interestingly, we uncover a strong correlation of the model parameter of the basal MET phosphorylation rate and the clinical outcome upon liver surgery. Since the extent of basal MET phosphorylation is a similarly good predictor of the clinical outcome as the estimated rate, the quantification of MET phosphorylation in surgical liver samples might provide a readily accessible readout to predict the clinical outcome. Taken together, patient-specific pMET levels could be exploited to assess the health status of the liver and to estimate the risk of a patient to suffer from liver failure after surgery.

## Materials and Methods

### Mouse housing and feeding

Standard diet control mice: Male C57BL/6N mice (Charles River) control mice were housed at the German Cancer Research Center (DKFZ) animal facility under a constant light/dark cycle and allowed ad libitum access to water and food and maintained on a standard mouse diet (KLIBA NAFAG 3437). The experiments were approved by the governmental review committee on animal care of the state Baden-Württemberg, Germany (reference number G- 14/17 and G-33/17).

Western diet-fed mice: Male C57BL/6N mice (Charles River) of 8 weeks of age were housed at the Leibniz-Institut für Arbeitsforschung an der Technischen Universität Dortmund (IfaDo) under a constant light/dark cycle and allowed ad libitum access to water and food and maintained on a Western diet (Research Diet Inc., D09100301, including trans fats) containing 40% kcal fat, 40% kcal carbohydrates and 2% weight cholesterol for 12 or 13 weeks. The experiments were approved by the governmental review committee of animal care of the state Nordrhein-Westfalen, Germany (reference number 84-02.04.2017.A177).

Transgenic R26p-Fucci2 mice: R26p-Fucci2 mice were obtained from RIKEN Center for Developmental Biology (CDB) and recovered by embryo transfer. The mice were housed at the German Cancer Research Center (DKFZ) animal facility under a constant light/dark cycle and allowed ad libitum access to water and food. Heterozygous R26p-Fucci2 mice were bred with wild type C57BL/6N mice to maintain the line. Only male animals were used for the performed analyses. Animals were either maintained on a standard mouse diet (KLIBA NAFAG 3437) or switched to a Western diet (WD, Research Diet Inc., D16022301, no trans fats) at the age of 8 weeks and fed with WD for 12 weeks. The experiments were approved by the governmental review committee of animal care of the state Baden-Württemberg, Germany (reference number A24/10, G-14/17, G-33/17).

### Isolation of primary mouse hepatocytes

Mice of final age of 20 to 21 weeks were used for primary mouse hepatocyte isolation. Hepatocytes were isolated according to a standardized procedure with adaptations to improve yields for WD-fed mice (Mueller *et al*., 2015). Anesthesia was carried out by intraperitoneal injection of 11.25 mg per 100 mg body weight ketamine hydrochloride (100 mg/ml, zoetis), 1.6 mg per 100 mg body weight xylazine hydrochloride (2% (w/v), Bayer HealthCare) and 1.5 mg per 100 mg acepromazine (cp-pharma). The abdominal cavity was opened and the vena cava inferior or portal vein was cannulated with a 24G venous catheter to enable perfusion of the liver. The liver was perfused with EGTA-containing buffer (0.6 % (w/v) glucose, 105 mM NaCl, 2.4 mM KCl, 1.2 mM KH_2_PO_4_, 26 mM Hepes, 490 M L-glutamine (Gibco), 512 μM EGTA, 15 % (v/v) amino acid solution (270 mg/l L-alanine, 140 mg/l L-aspartic acid, 400 mg/l L-asparagine, 270 mg/l L-citrulline, 140 g/l L-cysteine hydrochloride monohydrate, 1 g/l L-histidine monohydrochloride monohydrate, 1 g/l L-glutamic acid, 1 g/l L-glycine, 400 mg/l L-isoleucine, 800 mg/l L-leucine, 1.3 g/l L-lysine monohydrochloride, 550 mg/l L-methionine, 650 mg/l L-ornithine monohydrochloride, 550 mg/l L-phenylalanine, 550 mg/l L-proline, 650 mg/l L-serine, 1.35 g/l L-threonine, 650 mg/l L-tryptophane, 550 mg/l L-tyrosine, and 800 mg/l L-valine; pH 7.6) ; pH 8.3) for 5 min and collagenase-containing buffer (0.6 % (w/v) glucose, 105 mM NaCl, 2.3 mM KCl, 1.2 mM KH_2_PO_4_, 25 mM Hepes, 490 μM L-glutamine (Gibco), 5.3 mM CaCl_2_, 12 % (v/v) amino acid solution, 444 μg/ml collagenase type 1-A; pH 8.3) for up to 10 min at a flow rate of 8 ml/min. The portal vein or vena cava inferior was incised to allow sufficient buffer outflow. Following perfusion, the liver was withdrawn and transferred into suspension buffer (0.6 % (w/v) glucose, 105 mM NaCl, 2.4 mM KCl, 1.2 mM KH_2_PO_4_, 26 mM Hepes, 1 mM CaCl_2_, 0.4 mM MgSO_4_, 0.2 % (w/v) BSA, 490 μM L-glutamine (Gibco), 15 % (v/v) amino acid solution; pH 7.6). Hepatocytes were isolated by disrupting the liver capsule and filtering the resulting cell suspension through a 100 µm cell strainer. Cells were washed by centrifugation at 50 × g for 5 min at 4°C or twice at 50 × g for 2 min at room temperature and resuspended in adhesion medium (phenol red-free Williams E medium (PAN Biotech) supplemented with 10 % (v/v) FCS (Gibco), 0.1 μM dexamethasone, 0.1 % (v/v) insulin, 2 mM L-glutamine, 1 % (v/v) penicillin/streptomycin (Gibco)). Cell yield and vitality were determined by Trypan Blue staining using a Neubauer counting chamber. Preparations with vitality greater than 70% were employed for experiments. For each experiment cells from a single mouse were used, or if necessary, cells were pooled from up to three mice.

### Collection of patient tissue samples

Primary human hepatocytes were isolated from samples taken from the specimen of patients undergoing major hepatectomy at the Department of General, Visceral, and Transplantation Surgery of Heidelberg University Hospital. All patients were screened for eligibility irrespective of the indication for major hepatectomy and included provided they have signed the valid informed consent form. Informed consent of the patients for the use of tissue for research purposes was obtained corresponding to the ethical guidelines of University Hospital Heidelberg (reference number S-557/2017). A non-tumor tissue sample with intact Glisson’s capsule, weighing approximately 20 g was collected and immediately transported in William’s E medium (PAN Biotech) to the laboratory for further processing.

### Isolation of primary human hepatocytes

Cell isolation took place under a fume hood in sterile conditions using a standardized protocol adapted from Kegel *et al*. (2016). 3 to 8 cannulas were placed in the vessels of the non-capsuled surface of the liver sample. The cannulas were fixed with Histoacryl tissue glow and the blood was flushed from the tissue. The liver tissue was perfused with 500 ml 39°C 1× Perfusion Solution (142 mM NaCl, 6.7 mM KCl, 10 mM HEPES 12.5 mM EGTA, 6.25 mM N-Acetyl-L-Cystein) using a flow rate of 60 to 70% for 20 to 30 min until the liver tissue became light yellow. Afterwards the perfusion fluid was changed to 39°C digestion solution (33.5 mM NaCl, 3.35 mM KCl, 50 mM HEPES, 0.25 % BSA, 10 % FCS, 1 mg/ml Collagenase P) for 15 min. To stop the digestion the liver sample was rinsed with ice-cold Stop Solution (20 % FCS in DPBS (PAN Biotech)). After removing the cannulas, a scalpel was used to open the liver tissue. By flushing with Stop Solution and shaking the tissue gently, the cells were released from the tissue. The cell suspension was collected and filtered through a 100 µm cell strainer to 50 ml plastic falcons. The cell suspension was centrifuged at 50 × g, 5 min, 4°C. The cell pellet was washed with DPBS (PAN Biotech) followed by another round of centrifugation. Afterwards, the pellet was resuspended in adhesion medium. Cell yield and vitality were determined by Trypan Blue staining using a Neubauer counting chamber.

### Collection of human blood samples and plasma extraction

The blood of the patients was taken at the University Hospital Heidelberg one day before and 1, 3 and 7 days after surgery. Informed consent of the patients for the use of blood for research purposes was obtained corresponding to the ethical guidelines of University Hospital Heidelberg (reference number S-557/2017). For collecting blood, a vein in the arm bend was punctured, after a two-time disinfection with an alcoholic skin antiseptic. The blood was collected in EDTA 2.7 ml (Sarstedt). To receive the blood plasma the blood samples were centrifuged at full speed briefly and the supernatant was collected and frozen in -80°C until further use.

### Analysis of patient plasma cytokines

To analyze the preoperatively and postoperatively collected blood plasma samples of the patients, Bio-Plex Pro Cytokine, Chemokine, and Growth Factor Assay (Biorad) and the Bio-Plex Pro TGFβ Assay (Biorad) was used for multiplexing according to the standardized protocol established by Biorad. For multiplexing, HGF, IL6, IL8, PDGF, TGFβ1 and TNFα were analyzed. The blood plasma was thawed on ice and centrifuged twice (1000 × g, 15 min at 4°C). The samples were diluted 1:4 using Bio-Plex sample diluent. 50 µl coupled beads mix were pipetted to each well of the assay plate after vortexing for 30 s at medium speed. Then the beads were washed twice with the Bio-Plex wash buffer. After vortexing, 50 µl of the pre-diluted standards and samples were added. Samples and standards were assayed in technical duplicates. The plate was covered with sealing tape to protect from light and incubated on a shaker for 30 min at 850 ± 50 rpm. After the incubation, the beats were washed three times with washing buffer before 25 µl of the detection antibody mix was added. The plate was covered with sealing tape and incubated for a second time on the shaker for 30 min at 850 ± 50 rpm at room temperature. The incubation was followed by three more washing steps and the addition of 50 µl SA-PE, which was incubated for 10 min at 850 rpm ± 50 rpm for 10 min. After the final three washing steps the beads were re- suspended in 125 µl assay buffer and were shaken at 850 ± 50rpm for 30 s at room temperature and then measured with a Bio-Plex 200 reader.

For the TGFβ kit, after a 15 min centrifugation at 1000 × g and 10 min at 10 000 × g both at 4°C, acid (1 M HCl) and samples were added in 1:5 proportions and incubated at room temperature for 10 min. The samples were neutralized by adding 1 volume of base (1.2 M NaOH / 0.5 M HEPES) and vortexing. The (untreated) sample was in total diluted 1:16 with Bio-Plex sample diluent. After adding 50 µl of magnetic beads, washing two times, and pipetting 50 µl of standards and samples the plate was covered with sealing tape and incubated on a shaker at 850 ± 50 rpm for 2 h at room temperature. The incubation was followed by three more washing steps and the addition of 25 µl antibodies, which were incubated for 1 h at 850 ± 50 rpm at room temperature. The samples were washed three times and incubated with 50 µl of SA-PE for 30 min at 850 ± 50 rpm at room temperature. After washing three times again the beads were re-suspended in 125 µl assay buffer at each well and incubated for 30 s at 850 ± 50 rpm before the plate was measured with a Bio-Plex 200 reader.

### Collection of patient characteristics

Patient data (Dataset EV5) were prospectively collected by extraction from the electronic patient record system of the Heidelberg University Hospital. The collected information included clinic demographic data, preoperative clinical course, laboratory values, intraoperative variables, complications, time to event data, diagnose-related information, pre- operative treatment and histopathology reports. From this data, the following scores were obtained: Clavien Dindo score (Dindo *et al*., 2004), Charlson comorbidity index (Charlson *et al*, 1987) and comprehensive complication index (Slankamenac *et al*., 2013).

### Cultivation and stimulation of primary hepatocytes

Isolated primary hepatocytes were plated on collagen I-coated cell ware in adhesion medium. Using primary mouse hepatocytes, for the dose response and time course experiments 2 × 10^6^ cells were seeded per 6 cm dish. For live cell imaging 7.5 × 10^3^ primary mouse hepatocytes were seeded per well of a 96-well plate. For time course experiments with primary human hepatocytes, 2.5 × 10^6^ cells were seeded per 6 cm dish. For proliferation experiments, 150 000 cells per well were seeded on 6-well plates. Following plating, cells were allowed to adhere for 4 h (SD mouse hepatocytes) or 4 to 6.5 h (WD mouse hepatocytes and human hepatocytes). Subsequently, hepatocytes were washed, twice vigorously and once gently, with PBS (PAN Biotech) to remove unattached cells and cultured in serum-free medium overnight. During all incubation times, the cells were cultured at 37°C, 5% CO_2_ and 95% relative humidity.

For dose response and time course experiments, cells were washed gently three times and supplied with serum- and dexamethasone-free medium for 6 h. Primary mouse hepatocytes were stimulated with 0.1, 1, 2, 4, 10, 20, 40, 80, 100 and 120 ng/ml of recombinant mouse HGF (rmHGF, R&D systems) for 10 min for dose response experiments and with 40 ng/ml HGF for 5, 10, 20, 40, 60, 120, 180 and 240 min for time course experiments. Unstimulated control plates were taken out of the incubator together with stimulated plates and a zero dose / time point was taken in addition.

Primary human hepatocytes were stimulated with 40 ng/ml recombinant human HGF (R&D systems) for 1, 3, 5, 10, 20, 40, 60 and 120 min. Unstimulated control plates were taken out of the incubator together with stimulated plates and a zero time point was taken in addition.

### Live cell imaging of the cell cycle progression in primary mouse hepatocytes

Seeded primary mouse hepatocytes from R26p-Fucci2 transgenic mice were supplemented with purified AAV encoding Histone2B-mCerulean produced using a triple transfection protocol by Dirk Grimm (Heidelberg University) (Grimm, 2002) for the duration of cell adhesion. Following adhesion, cells were washed twice with serum-free medium and incubated in serum-free medium for 24 h before stimulationShortly before time-lapse microscopy, media were changed to 100 µl fresh serum-free medium supplemented with 40 ng/ml HGF. Hepatocytes were imaged with a Nikon Eclipse Ti Fluorescence microscope controlled with NIS- Elements software. Temperature (37°C), CO_2_ (5%) and humidity were held constant by an incubation chamber enclosing the microscope and a 96-well plate stage insert. Four channels were acquired: brightfield channel, CFP channel (Histone2B- mCerulean), RFP channel (mCherry-hCdt1), and YFP channel (mVenus-hGeminin). Time lapse microscopy was performed for up to 65 h with a sampling rate of 20 min.

Image analysis was performed using the Fiji software. Background subtraction was performed with rolling ball method. CFP channel was used for segmentation of nuclei using a standard threshold-based algorithm. Mean RFP and YFP intensity was quantified for all segmented nuclei at each time frame.

### Snapshot population analysis of Fucci2 data

The programming language R was used to perform subpopulation analysis of Fucci2 data. All segmented nuclei within a specified time window between 42 and 54 h were displayed as a scatter plot to quantify percentage of subpopulations in a given cell cycle phase. Four subpopulations were defined using arbitrary thresholds for mCherry and mVenus signals based on unstimulated condition as a reference. The four compartments were defined as RFPhigh/YFPlow (G1-phase, red), RFPhigh/YFPhigh (S-phase, orange), RFPlow/YFPhigh (G2-phase, green), RFPlow/YFPlow (M-phase, gray).

### Time-resolved analysis of cell cycle progression in single cells

Manual segmentation and tracking were performed to obtain single cell tracks. To perform reliable quantification of cell cycle duration and cell cycle entries, only cells that were present in the field of view for the whole duration of the time lapse were considered. A region of interest (ROI) was defined in individual nuclei in the image of the CFP channel (Histone2B- mCerulean) for each frame. At the event of cell division, only one of the daughter cells was followed until the end of the whole time-lapse experiment. The obtained set of ROIs was applied to the RFP (mCherry-hCdt1) and YFP (mVenus-hGeminin) channels to extract the RFP and YFP mean intensity values after background subtraction by rolling ball method. The two thresholds for the RFP and YFP channels defined in the snapshot analysis were applied to assign cells to the four cell cycle phases, represented as the respective color in the heatmap. To account for fluctuations in the expression of the Fucci2 fluorescent probes, cell cycle entries were considered as transition from S to G2-phase.

### Cell lysis

Cells were lysed in 500 µl (mouse hepatocytes) or 300 µl (human hepatocytes) RIPA buffer (50 mM Tris pH 7.4, 150 mM NaCl, 1 mM EDTA, 1 mg/ml Deoxycholic acid, 0.5 mM Na3VO4, 2.5 mM NaF, 1% NP40, 0.1% AEBSF, 0.1% AP) on ice, incubated while rotating at 4°C for 20 min and were sonicated for 30 s (Amplitude: 80%, 0.1 on, 0.5 off) (human hepatocytes only) followed by centrifugation at 4°C for 10 min at 20 817 × g. The supernatant representing protein lysates were stored at -80°C. Pierce BCA Protein Assay Kit (Thermo Fisher) was used to determine the protein concentration in the total cell lysates.

### Quantitative immunoblotting

For total cell lysate analysis 30 µg protein (mouse hepatocytes) or 20 to 30 µg (human hepatocytes) was filled up to 25 µl total volume with RIPA buffer, mixed with 25 µl 2× sample buffer (4% SDS, 100 mM tris-HCl pH 7.4, 20% glycerol, 200 mM DTT, bromophenol blue, 10% β-mercaptoethanol), incubated for 3 to 5 min at 95°C and used for quantitative immunoblotting. Total cell lysates and immunoprecipitation samples were loaded on 10% polyacrylamide gels in a randomized order to avoid correlated blotting errors (Schilling *et al*, 2005). Magic Marker (Invitrogen) and Precision Plus Protein Standard (Biorad) were loaded in addition. The proteins were separated by discontinuous gel electrophoresis (SDS-PAGE) (40 mA per gel for 3 h) covered with Laemmli Buffer (192 mM glycin, 25 mM Tris, 0.1% SDS). The separated proteins were transferred in a semi-dry system onto PVDF membranes (Immobilon P, Merck Millipore) using transfer buffer (192 mM glycine, 25 mM Tris, 0.075\% SDS, 0.5 mM Na3VO4, 15% EtOH) (Hoefer TE77 semi-dry transfer system (250 mA per blot for 1h)). The membrane was blocked by drying after being soaked in ethanol followed by a 10 min reactivation in TBS-T (10 mM Tris pH 7.4, 150 mM NaCl, 0.2% Tween-20). The membranes were incubated with primary antibodies (against MET (Sanza Cruz and Cell Signaling), pMET Tyr 1234/1235, pAKT Thr 308, pAKT Ser 473, AKT, pERK Thr 202/Tyr 204, ERK, pS6K Thr 389, S6K, pS6 Ser 235/236, S6 (all Cell Signaling)) in 2.5% BSA in TBS-T overnight at 4°C.

The membranes were washed twice with TBS-T for 5 min followed by a 1 h incubation with secondary Antibody (HRP Goat Anti-Rabbit (Dharmacon) or HRP Goat Anti-Mouse (Dharmacon), 1:10 000 in 2.5% BSA in TBS-T). After two additional washing steps with TBS-T and one with TBS the blots were developed by incubation with Amersham ECL Western Blotting Detection Reagents (GE Healthcare) for 2 min and detected on an ImageQuant (GE Healthcare). The proteins were quantified using ImageQuant TL software (GE Healthcare, version 7.0). For further detection, the blots were treated with stripping buffer (62.5 mM Tris pH 6.8, 2% SDS, 0.7% β-mercaptoethanol) for 20 min at 65°C to remove the bound antibodies. After washing with double-distilled water, blocking by drying, and reactivation by shaking in TBS-T, membranes were employed for further antibody binding. Alternatively, the secondary antibody was inactivated using 30% H_2_O_2_ for 15 min at 37°C followed by three washing steps in double-distilled water for 5 min each and reactivation in TBS-T.

### Proliferation assay

At time point 0 h, primary human hepatocytes were washed with PBS twice. For proliferation, primary human hepatocytes were stimulated by adding new serum and dexamethasone free medium containing 40 ng/ml recombinant human HGF. Medium without HGF was used for control cells. After 48 h the hepatocytes were washed once with PBS and the plates were frozen at -20°C. Control cells were washed with DPBS twice at time point 0 h and frozen at - 20°C. For DNA content measurement, staining with SybrGreen I (Invitrogen, S7563) was performed. The frozen cells were treated with 2 ml SybrGreen I working solution (PBS, 1:100 Triton X 100, 1:2 500 SybrGreen I) per well, incubated protected from light for 1 h. Signal emission was measured at 485 nm excitation using a Tecan Infinite Pro 200 reader. The optimal gain was determined with the plate for which the highest signal was expected and this value was used for all further measurements.

### Proteomics sample preparation

Lysates of unstimulated primary mouse hepatocytes were used for proteome analysis. Protein concentrations were determined by employing the BCA Protein Assay Kit (Pierce,Thermo Fisher). In total, 20µg of protein per sample was used for further processing. The disulfide bonds of proteins were reduced with 40mM Tris(2-carboxyethyl) phosphine (TCEP) and then alkylated with 10mM chloroacetamide (CAA) for 60min at 37°C. Protein digestion and clean-up were performed using an adapted version of the automated paramagnetic bead-based single-pot, solid-phase-enhanced sample-preparation (Auto-SP3) protocol (Muller *et al*, 2020) on the Bravo liquid handling platform (Agilent). Briefly, for bead preparation, Sera-Mag Speed Beads A and B (Ge Healthcare) were vortexed until the pellet was dissolved. The suspension was placed on a magnetic rack, and after one minute, the supernatant was removed. The beads were taken off the magnetic rack and suspended in water. This procedure was repeated three times. A total of 10µL of bead A was combined with 10µL of bead B, and the final volume was corrected to 100µL with H_2_O. To each sample, a total of 5µL of A+B beads were added. To induce the binding of the proteins to the beads, ethanol was added to each sample to a final 50% concentration (v/v). Samples were then incubated for 15 min at room temperature and 800 rpm. After the incubation step, samples were placed again on a magnetic rack, and after one minute, the supernatant was removed. Samples were taken off the magnetic rack and suspended in 80 % ethanol. This procedure was repeated three times. Finally, samples were reconstituted in 100mM TEAB buffer containing trypsin (enzyme/protein ratio of 1:25) and digested overnight on a shaker at 37 °C and 1000 rpm. After digestion, the recovered peptides were dried by vacuum centrifugation and stored at -80°C until use.

### LC-MS/MS Analysis

Nano-flow LC-MS/MS was performed by coupling an Ultimate 3000 HPLC (Thermo Fisher Scientific, USA) to a Orbitrap Exploris mass spectrometer (Thermo Fisher Scientific, Germany). Peptide samples were dissolved in 15 µl loading buffer (0.1% formic acid (FA), 2% ACN in MS-compatible H2O), and 2 µL were injected for each analysis. The samples were loaded onto a pre-column (trap) at higher flowrates (PEPMAP 100 C18 5µm 0.3mmx5mm, Thermo Scientific), using a loading pump and then a valve was switched to delivered to an analytical column (75 µm × 30 cm, packed in-house with Reprosil-Pur 120 C18-AQ, 1.9 µm resin, Dr. Maisch) at a flow rate of 3 µL/min in 98% buffer A (0.1% FA in MS-compatible H2O). After loading, peptides were separated using a 120 min gradient from 2% to 38% of buffer B (0.1% FA, 80% ACN in MS-compatible H2O) at 350 nL/min flow rate. The Orbitrap Exploris 480 was operated in data-independent (DIA) mode, with a m/z range of 350–1400. Full scan spectra were acquired in the Orbitrap at 120 000 resolution after accumulation to the set target value of 300% (100% = 1e6) and maximum injection time of 45 ms. The full scans were followed by DIA scans. A total of 47 isolation windows were defined, with a m/z range of 406-986. DIA scan spectra were acquired at 30 000 resolution after accumulation to the set target value of 1000% (100% =1e5) and maximum injection time of 54 ms. Normalized collision energy (NCE) was set to 28%.

### Database search and Data Analysis

All DIA–MS data files were analyzed with a directDIA workflow using Spectronaut 15.5 (Biognosys, Zurich, Switzerland). For the Pulsar search, the UniProt human-reviewed canonical reference proteome (20,593 entries, UP000005640 downloaded on December 7th, 2022) and the UniProt mouse-reviewed canonical reference proteome (20,404 entries, downloaded on December 7th, 2022) were used. The default settings for database match include: full specificity trypsin digestion, peptide length of between 7 and 52 amino acids and maximum missed cleavage of 2. N-terminal methionine was removed during preprocessing of the protein database. Carbamidomethylation at cysteine was used as a fixed modification, protein N-terminal acetylation and methionine oxidation were set as variable modifications. The false discovery rates (FDRs) were set as 0.01 for the peptide-spectrum match (PSM), peptide and protein identification. Other Spectronaut parameters for identification included maximum intensity as MZ extraction strategy, 0.01 precursor Qvalue Cutoff, 0.2 Precursor PEP Cutoff, stripped sequence as single hit definition. For quantification, identified (Qvalue) was set for precursor filtering, MaxLFQ as protein LFQ method and MS2 quantification with area as quantity type. For further analysis, only those proteins without missing values were considered. For the analysis of the proteome data the public server at usegalaxy.org was used (Afgan *et al*, 2016). Analysis was based on the R package *limma* (Ritchie *et al*., 2015). Statistical testing of significance was performed using the *voom* function from Bioconductor (Law *et al*, 2014). Significance was considered with adjusted p-value < 0.05. Functional annotation of canonical pathways was performed with the use of Ingenuity Pathway Analysis (Krämer *et al*., 2014) by QIAGEN. Thresholds were set to an adjusted p-value of 0.05 and a log_2_ fold change < -0.5 or > 0.5.

### Data processing and estimation of uncertainties

Immunoblot data from individual experiments were scaled and thereby aligned for each target using the R package *blotIt* (Kemmer *et al*, 2022). Uncertainties, corresponding to 1*σ* confidence intervals, were estimated along with scaling factors for each experiment using the scaling model 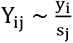 with measurements *Y_ij_*, scaling factors *s_j_* and scaled values *y_i_*, combined with a relative error model 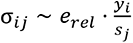 with error parameter *e_rel_* and estimated errors *σ_ij_*. The biological effects *i* were distinguished for individual targets, measurements time points, mouse diets and HGF stimulation doses. Scaling effects *j* were distinguished per experiment and target. For the mouse model 4 466 raw data points were aligned by *blotIt* to obtain 772 data points with confidence intervals that served for calibration of the mouse model. In addition, 209 raw data points for protein abundances obtained from DIA measurements were log_2_ transformed and averaged to obtain 22 data points with calculated standard deviations. Finally, the number of molecules per cell of AKT determined by quantitative immunoblotting was also utilized for model calibration. For patient-derived primary human hepatocytes, 2443 raw data points were aligned by *blotIt* to obtain 1106 data points with confidence intervals that served for calibration of the human model. In addition, 112 raw data points for protein abundances obtained from DIA measurements were log_2_ transformed and averaged to obtain 56 data points with calculated standard deviations.

### Parameter estimation and model development for primary mouse hepatocytes

The mathematical modeling was performed in the R package *dMod*, which provides an environment for the development of ODE models, parameter estimation and uncertainty analysis (Kaschek *et al*, 2019). The final ODE model was composed of 23 species and 26 reactions derived from the law of mass-action (Dataset EV1). Measurements were mapped to model states by means of observables as displayed in Dataset EV2. The estimated parameter set corresponding to the global optimum along with profile-likelihood based confidence intervals is listed in Dataset EV3. Log-transformation of parameters was used to ensure positivity and numerical stability. A set of parameter transformations (Dataset EV4) was used to incorporate the calculation of analytical steady-state expressions (Rosenblatt *et al*, 2016) and reformulations necessary for model reduction (Maiwald *et al*, 2016). Parameter values were estimated using the maximum likelihood method, performing a deterministic multi-start optimization with the trust region optimizer (R package *trust*) and starting from 250 randomly chosen parameter sets. Results of the optimization run for the final model are displayed as waterfall plots (Raue *et al*, 2013) in Fig EV1D. The global optimum was found in 164 out of the 250 starts for the final model. The profile likelihood (Raue *et al*., 2009) was used to assess the identifiability of parameters and determine confidence intervals for the estimated values of 22 initial protein concentrations, 13 scaling and offset parameters, and 27 dynamical parameters of the mouse model. For the identification of mechanistic differences in the signal transduction of SD and WD hepatocytes models differing in their number and the selection of diet-specific parameters were compared using the Bayesian information criterion (Schwarz, 1978).

### Application of the mouse model to primary human hepatocytes

The calibrated mouse model was used for the analysis of measurements derived from primary human hepatocyte. The parameter values were fixed to the estimates obtained for the mouse data except for a parameter subset that was identified by model comparison based on the Bayesian information criterion. These human-specific parameters comprise the basal MET phosphorylation rate and protein abundances, which were estimated as patient-specific, scaling and offset parameters, as well as the HGF-induced MET phosphorylation rate and the pMET degradation rate, which were implemented as human-specific parameters identical for all patients. These non-fixed parameters were re-estimated using the maximum likelihood method. The resulting parameter estimates are summarized in Dataset EV6.

## Data availability

The mass spectrometry proteomics data have been deposited to the ProteomeXchange Consortium through the PRIDE partner repository (Vizcaino *et al*, 2014) with the data set identifier PXD043007 (human) Reviewer account details:

**Password:** xrjgNBIk and PXD041563 (mouse) Reviewer account details:

**Password:** ifue4xbD

The ODE models developed for the mouse and the human setting are available on BioModels (Malik-Sheriff *et al*, 2020) in SBML and PEtab (Schmiester *et al*, 2021) formats under the identifier MODEL2306280002. Reviewer account details:

**Username:** reviewerForMODEL2306280002

**Password:** J8CY0Z

## Acknowledgements

The authors thank the Proteomics Core Facility of the German Cancer Research Center (DKFZ), especially Dominic Helm and Luisa E. Schwarzmüller, for providing support in establishing the DIA workflow and data analysis. Furthermore, we would like to thank Dirk Grimm for production of AAV particles and Marcus Rosenblatt for fruitful discussions during the development of the mathematical model. We thank Katharina Belgasmi, Lena Vlasov, Sandra Bonefas and Alexander Held for technical assistance. We thank the Nikon Imaging Center (Heidelberg University) for providing access to their facility. This work was supported by the German Ministry of Education and Research (BMBF) within the LiSyM network [031L0042, 031L0045, 031L0048, 031L0049, 031L0052] and the LiSyM-Cancer networks SMART-NAFLD [031L0256A, 031L0256B, 031L0256G], C-TIP-HCC [031L0257C, 031L0257D, 031L0257K], the MSCoreSys network SMART-CARE [031L0212B] and by the German Center for Lung Research (DZL) [82DZL004A4]. The authors acknowledge support by the Open Access Publication Fund of the University of Freiburg and the state of Baden-Württemberg through bwHPC.

## Author contributions

Conceptualization: I.B., S.K., S.B., L.A.D., M.S., J.T., U.K.; methodology: S.K., B.H.; formal analysis: I.B., S.K., A.V.; investigation: I.B., Y.D.,A.V., S.F.; resources: K.H., A.G., J.G.H.; data curation: S.K.; writing---original draft preparation: I.B., S.K., S.B., M.S.; writing---review and editing: J.T., U.K.; visualization: I.B., S.K., S.B.; supervision: M.S., U.K., J.T.; project administration: U.K.; funding acquisition: U.K., J.T. All authors have read and agreed to the published version of the manuscript.

## Supporting Information

Dataset EV1: Model reactions

Dataset EV2: Model observation functions

Dataset EV3: Parameter values

Dataset EV4: Parameter transformations

Dataset EV5: Patient anamnesis

Dataset EV6: Model parameters of primary human hepatocytes

Dataset EV7: Clinical features

## Expanded View Figure legends

**Figure EV1.**
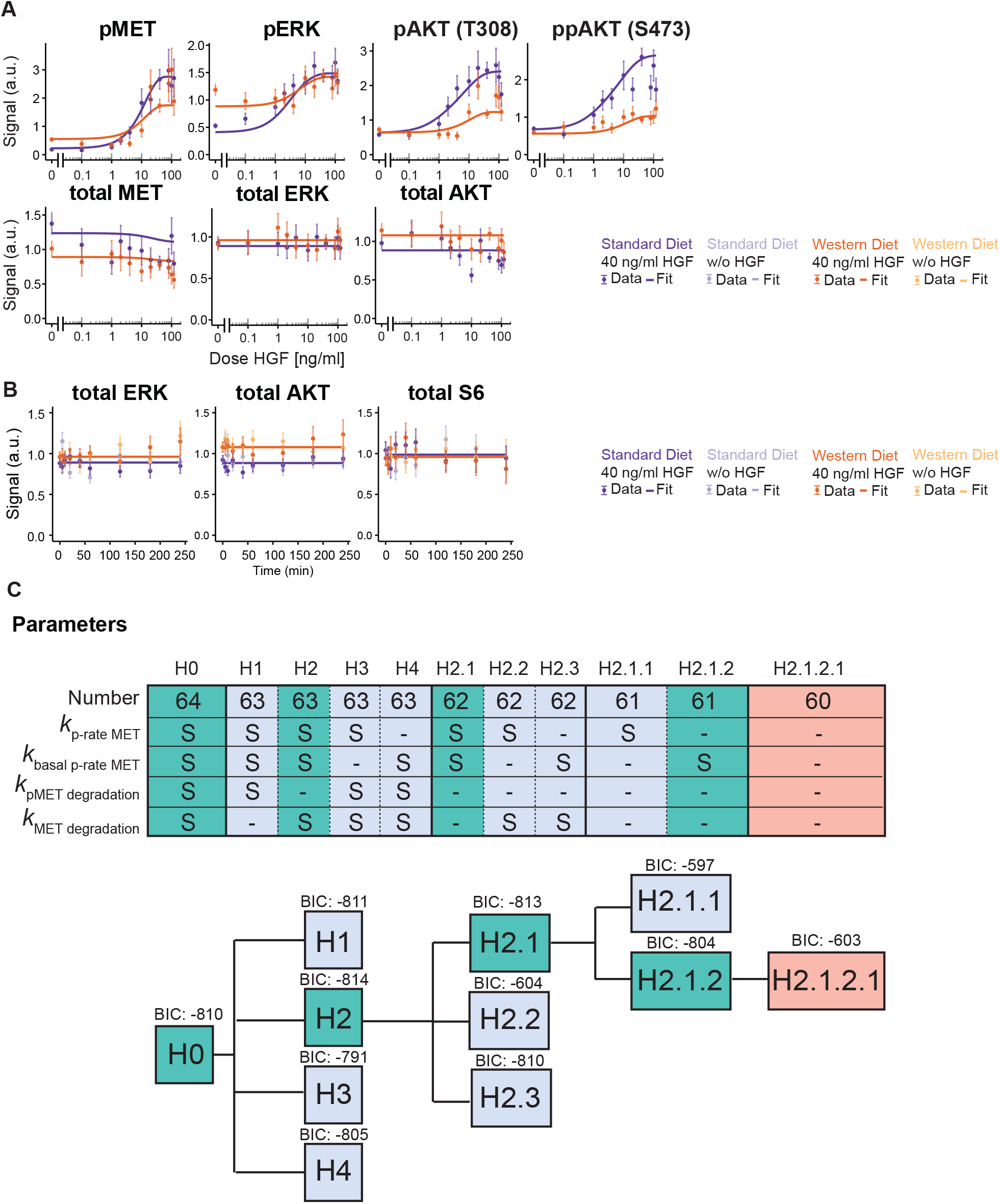
Quantitative data for calibration of the mouse model. A. Model calibration with HGF dose-resolved signal transduction measurements in primary hepatocytes of SD and WD mice. Hepatocytes were growth factor-depleted and stimulated with different doses of HGF. Lysates were subjected to quantitative immunoblotting. Measurements are represented by filled circles with errors representing 1σ confidence intervals estimated from biological replicates (N = 9) using a combined scaling and error model. Model trajectories are represented by solid lines. B. Model calibration with HGF time-resolved signal transduction measurements in primary hepatocytes of SD and WD mice. C. A Bayesian information criterion (BIC) analysis was performed to determine the diet specific parameters needed to describe the experimental data. The threshold for rejection was set to ΔBIC = 10 as suggested (Lorah & Womack, 2019). H0 including 64 parameters could be reduced to H2.1.2 including 61 parameters, suggesting that only the basal phosphorylation rate of the HGF receptor MET was dysregulated between diets.

**Figure EV2.**
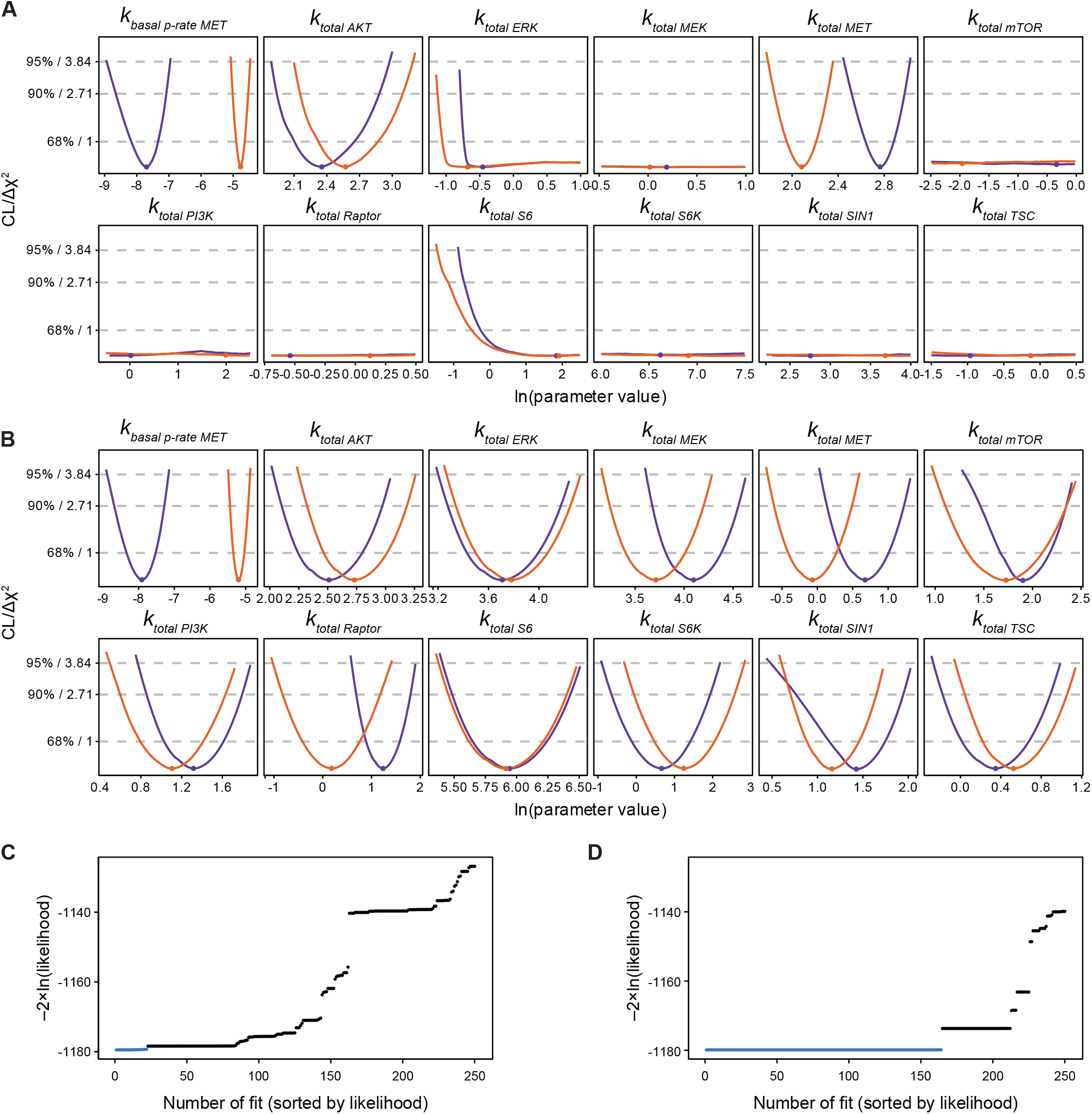
Impact of the DIA data on identifiability and convergence. A. The profile likelihood as a measure of parameter identifiability (Raue *et al*., 2009) is depicted for all dysregulated parameters before implementation of the DIA data. If the negative log likelihood reaches a statistical threshold in both directions, the parameter has defined confidence bounds and is therefore called identifiable. If this limit is not reached on both sides, the parameter is classified as unidentifiable. Solid lines indicate the profile likelihood of dysregulated parameters for SD (purple) and WD (orange) along with the optimal parameter values as dots. Dashed lines depict thresholds for the confidence interval assessment. B. The profile likelihood as a measure of parameter identifiability is depicted for all dysregulated parameters after implementation of the DIA data. C. The convergence of the optimization before implementation of the DIA data is assessed based on a waterfall plot (Raue *et al*., 2013). This plot depicts the results of 250 optimization runs starting from randomly selected parameter sets sorted by the negative log likelihood. The global optimum, indicated in blue, was reached in 22 of the 250 cases. D. The convergence of the optimization after implementation of the DIA data is assessed based on a waterfall plot. The global optimum, indicated in blue, was reached in 164 out of 250 cases.

**Figure EV3.**
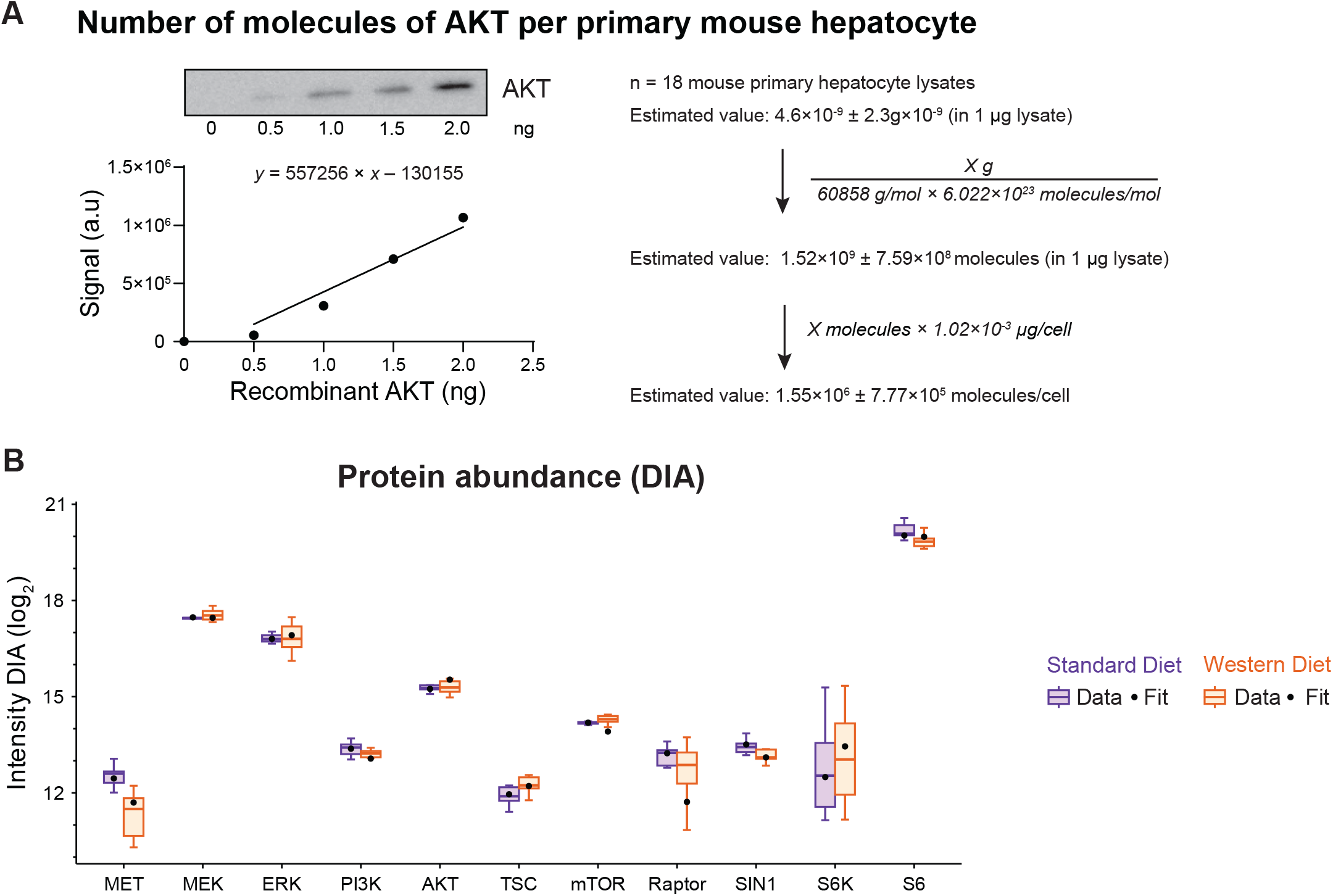
Absolute quantification of AKT and model-based estimations of total protein abundance. A. Absolute number of molecules of AKT per primary mouse hepatocyte was determined by quantitative immunoblotting. Based on a dilution curve of recombinant AKT, the number of molecules of AKT in 1 µg lysate was determined. This value was converted with the total protein content per primary mouse hepatocyte into the number of molecules of AKT per cell. B. Measurements for protein abundances, also from primary hepatocytes of SD and WD mice, were included as additional data for model calibration. Lysates of unstimulated hepatocytes were subjected to data-independent mass spectrometry analysis. Resulting data was normalized using label-free quantification and represented as boxplots. Model fits are represented as black dots.

**Figure EV4.**
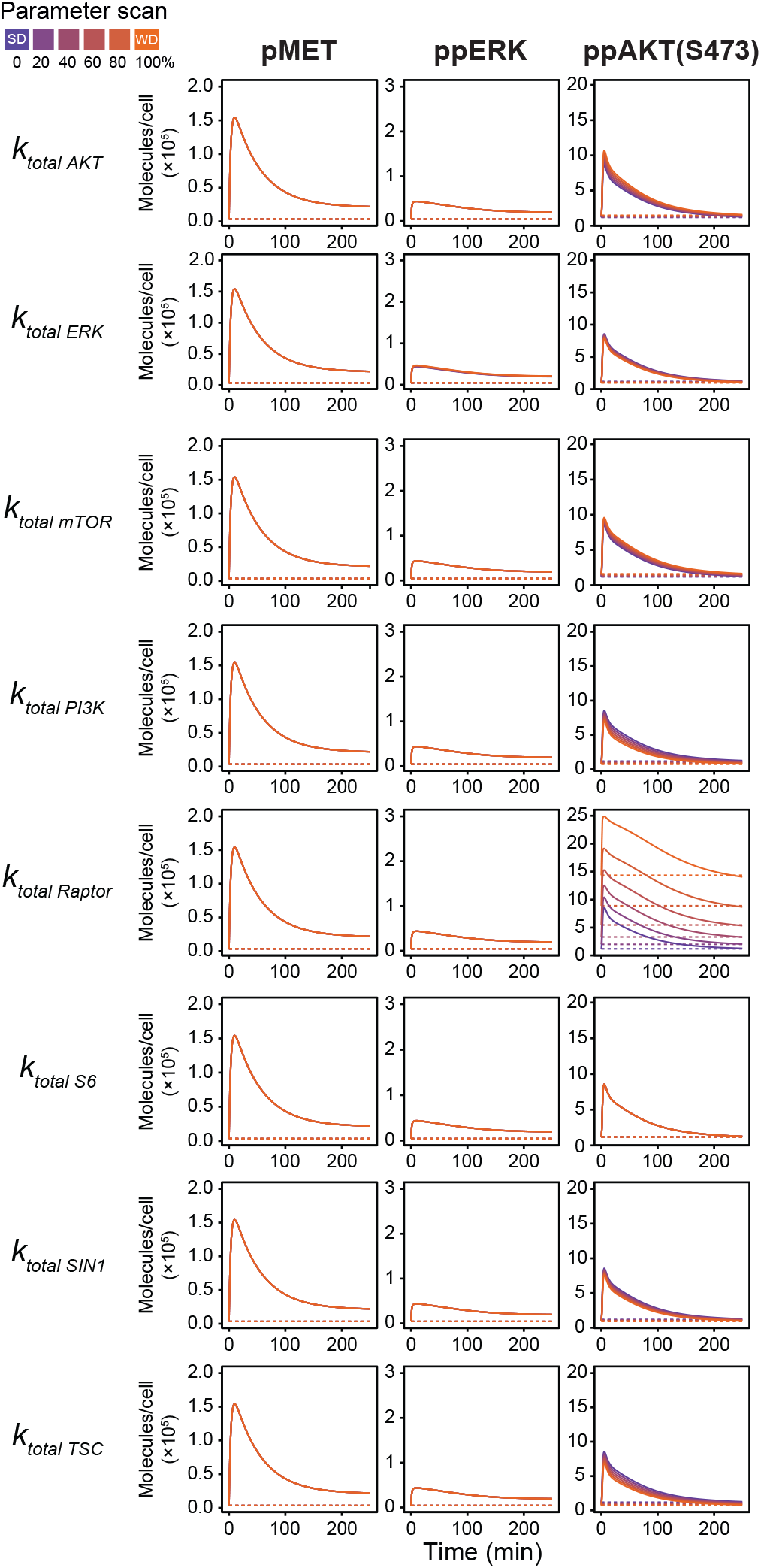
Influence of dysregulated parameters on protein dynamics. Influence of dysregulated parameters on protein dynamics. Phosphorylation dynamics of pMET, pERK and ppAKT were simulated based on the SD parameter set. The influence of each dysregulated parameter was analyzed by scanning its value from the estimate obtained for SD (purple) to the estimate obtained for WD (orange). Solid lines indicate model trajectories after HGF stimulation and dashed lines indicate the control condition

**Figure EV5.**
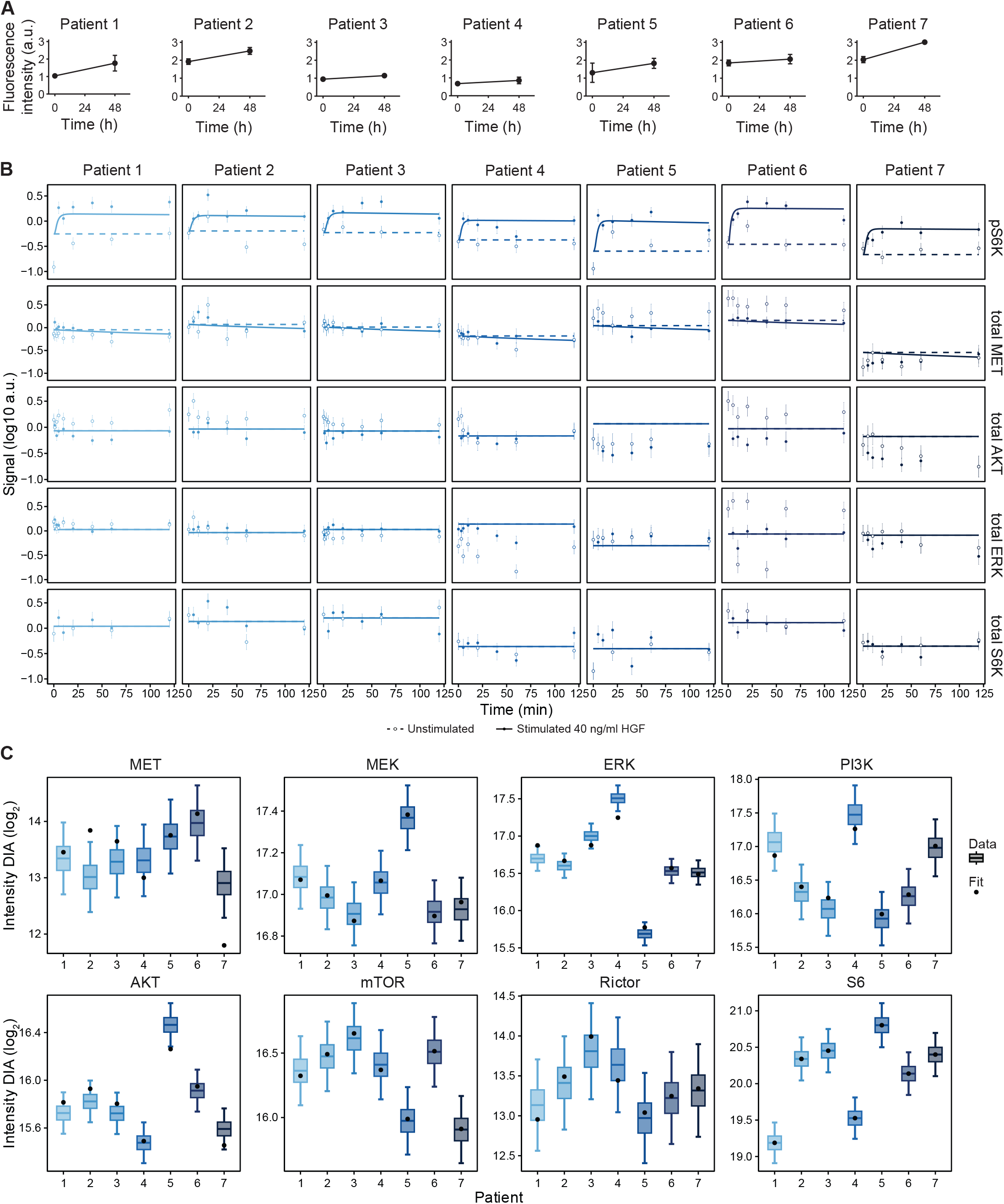
Quantitative data for calibration of the human model. A. Proliferation measurements in primary human hepatocytes. Primary human hepatocytes were isolated by liver perfusion, were allowed to adhere to 6-well plates and cultured in serum-free medium overnight. Cells were stimulated with 40 ng/ml HGF for 48 h. DNA content was measured by staining with SybrGreen I. B. Model calibration with time-resolved signal transduction measurements in primary human hepatocytes. Hepatocytes were growth factor-depleted and stimulated with HGF or left untreated. Lysates were subjected to quantitative immunoblotting. Measurements are represented by circles with errors representing 1*σ* confidence intervals estimated from technical replicates (n = 2 to 6) using a combined scaling and error model. Model trajectories are represented by solid lines. C. Measurements for protein abundances derived from primary patient hepatocytes were included as additional data for model calibration. Lysates of unstimulated hepatocytes were subjected to data-independent mass spectrometry analysis. Resulting data was normalized using label-freequantification and represented as boxplots. Model fits are represented as black dots.

## References

Abe T, Sakaue-Sawano A, Kiyonari H, Shioi G, Inoue K-i, Horiuchi T, Nakao K, Miyawaki A, Aizawa S, Fujimori T (2013) Visualization of cell cycle in mouse embryos with Fucci2 reporter directed by Rosa26 promoter. Development 140: 237–246

Adlung L, Kar S, Wagner MC, She B, Chakraborty S, Bao J, Lattermann S, Boerries M, Busch H, Wuchter P et al (2017) Protein abundance of AKT and ERK pathway components governs cell type-specific regulation of proliferation. Mol Syst Biol 13: 904

Afgan E, Baker D, van den Beek M, Blankenberg D, Bouvier D, Cech M, Chilton J, Clements D, Coraor N, Eberhard C et al (2016) The Galaxy platform for accessible, reproducible and collaborative biomedical analyses: 2016 update. Nucleic Acids Res 44: W3–W10

Allaire M, Gilgenkrantz H (2018) The impact of steatosis on liver regeneration. Horm Mol Biol Clin Investig 41

Bottaro DP, Rubin JS, Faletto DL, Chan AM, Kmiecik TE, Vande Woude GF, Aaronson SA (1991) Identification of the hepatocyte growth factor receptor as the c-met proto-oncogene product. Science 251: 802–804

Bussolino F, Di Renzo MF, Ziche M, Bocchietto E, Olivero M, Naldini L, Gaudino G, Tamagnone L, Coffer A, Comoglio PM (1992) Hepatocyte growth factor is a potent angiogenic factor which stimulates endothelial cell motility and growth. J Cell Biol 119: 629–641

Charlson ME, Pompei P, Ales KL, MacKenzie CR (1987) A new method of classifying prognostic comorbidity in longitudinal studies: development and validation. J Chronic Dis 40: 373–383

Chembazhi UV, Bangru S, Hernaez M, Kalsotra A (2021) Cellular plasticity balances the metabolic and proliferation dynamics of a regenerating liver. Genome Res 31: 576–591

D’Alessandro LA, Klingmuller U, Schilling M (2022) Deciphering signal transduction networks in the liver by mechanistic mathematical modelling. Biochem J 479: 1361–1374

D’Alessandro LA, Samaga R, Maiwald T, Rho SH, Bonefas S, Raue A, Iwamoto N, Kienast A, Waldow K, Meyer R et al (2015) Disentangling the Complexity of HGF Signaling by Combining Qualitative and Quantitative Modeling. PLoS Comput Biol 11: e1004192

Dalle Pezze P, Sonntag AG, Thien A, Prentzell MT, Godel M, Fischer S, Neumann-Haefelin E, Huber TB, Baumeister R, Shanley DP et al (2012) A dynamic network model of mTOR signaling reveals TSC-independent mTORC2 regulation. Sci Signal 5: ra25

DeAngelis T, Morrione A, Baserga R (2010) Mutual interaction and reciprocal down- regulation between c-met and insulin receptor substrate-1. J Cell Physiol 224: 658–663

Dehlke K, Krause L, Tyufekchieva S, Murtha-Lemekhova A, Mayer P, Vlasov A, Klingmuller U, Mueller NS, Hoffmann K (2022) Predicting liver regeneration following major resection. Sci Rep 12: 13396

Dindo D, Demartines N, Clavien PA (2004) Classification of surgical complications: a new proposal with evaluation in a cohort of 6336 patients and results of a survey. Ann Surg 240: 205–213

Ghanemi A, Yoshioka M, St-Amand J (2020) Regeneration during Obesity: An Impaired Homeostasis. Animals (Basel) 10

Greco D, Kotronen A, Westerbacka J, Puig O, Arkkila P, Kiviluoto T, Laitinen S, Kolak M, Fisher RM, Hamsten A et al (2008) Gene expression in human NAFLD. Am J Physiol Gastrointest Liver Physiol 294: G1281–1287

Grimm D (2002) Production methods for gene transfer vectors based on adeno-associated virus serotypes. Methods 28: 146–157

Hall Z, Chiarugi D, Charidemou E, Leslie J, Scott E, Pellegrinet L, Allison M, Mocciaro G, Anstee QM, Evan GI et al (2021) Lipid Remodeling in Hepatocyte Proliferation and Hepatocellular Carcinoma. Hepatology 73: 1028–1044

Huang DQ, El-Serag HB, Loomba R (2021) Global epidemiology of NAFLD-related HCC: trends, predictions, risk factors and prevention. Nat Rev Gastroenterol Hepatol 18: 223–238

Jeffers M, Taylor GA, Weidner KM, Omura S, Vande Woude GF (1997) Degradation of the Met tyrosine kinase receptor by the ubiquitin-proteasome pathway. Mol Cell Biol 17: 799–808

Jia G, Aroor AR, Martinez-Lemus LA, Sowers JR (2014) Overnutrition, mTOR signaling, and cardiovascular diseases. Am J Physiol Regul Integr Comp Physiol 307: R1198–1206

Kaschek D, Mader W, Fehling-Kaschek M, Rosenblatt M, Timmer J (2019) Dynamic Modeling, Parameter Estimation, and Uncertainty Analysis in R. J Stat Softw 88: 1–32

Kegel V, Deharde D, Pfeiffer E, Zeilinger K, Seehofer D, Damm G (2016) Protocol for Isolation of Primary Human Hepatocytes and Corresponding Major Populations of Non- parenchymal Liver Cells. J Vis Exp: e53069

Kemmer S, Bang S, Rosenblatt M, Timmer J, Kaschek D (2022) BlotIt-Optimal alignment of Western blot and qPCR experiments. PLoS One 17: e0264295

Kok F, Rosenblatt M, Teusel M, Nizharadze T, Goncalves Magalhaes V, Dachert C, Maiwald T, Vlasov A, Wasch M, Tyufekchieva S et al (2020) Disentangling molecular mechanisms regulating sensitization of interferon alpha signal transduction. Mol Syst Biol 16: e8955

Krämer A, Green J, Pollard J, Jr., Tugendreich S (2014) Causal analysis approaches in Ingenuity Pathway Analysis. Bioinformatics 30: 523–530

Kwiecinski M, Noetel A, Elfimova N, Trebicka J, Schievenbusch S, Strack I, Molnar L, von Brandenstein M, Tox U, Nischt R et al (2011) Hepatocyte growth factor (HGF) inhibits collagen I and IV synthesis in hepatic stellate cells by miRNA-29 induction. PLoS One 6: e24568

Law CW, Chen Y, Shi W, Smyth GK (2014) voom: Precision weights unlock linear model analysis tools for RNA-seq read counts. Genome Biol 15: R29

Le Novere N, Hucka M, Mi H, Moodie S, Schreiber F, Sorokin A, Demir E, Wegner K, Aladjem MI, Wimalaratne SM et al (2009) The Systems Biology Graphical Notation. Nat Biotechnol 27: 735–741

Lorah J, Womack A (2019) Value of sample size for computation of the Bayesian information criterion (BIC) in multilevel modeling. Behav Res Methods 51: 440–450

Maiwald T, Hass H, Steiert B, Vanlier J, Engesser R, Raue A, Kipkeew F, Bock HH, Kaschek D, Kreutz C et al (2016) Driving the Model to Its Limit: Profile Likelihood Based Model Reduction. PLoS One 11: e0162366

Malik-Sheriff RS, Glont M, Nguyen TVN, Tiwari K, Roberts MG, Xavier A, Vu MT, Men J, Maire M, Kananathan S et al (2020) BioModels-15 years of sharing computational models in life science. Nucleic Acids Res 48: D407–D415

Mueller S, Huard J, Waldow K, Huang X, D’Alessandro LA, Bohl S, Borner K, Grimm D, Klamt S, Klingmuller U et al (2015) T160-phosphorylated CDK2 defines threshold for HGF dependent proliferation in primary hepatocytes. Mol Syst Biol 11: 795

Muller T, Kalxdorf M, Longuespee R, Kazdal DN, Stenzinger A, Krijgsveld J (2020) Automated sample preparation with SP3 for low-input clinical proteomics. Mol Syst Biol 16: e9111

Murtha-Lemekhova A, Fuchs J, Ghamarnejad O, Nikdad M, Probst P, Hoffmann K (2021) Influence of cytokines, circulating markers and growth factors on liver regeneration and post- hepatectomy liver failure: a systematic review and meta-analysis. Sci Rep 11: 13739

Oe H, Kaido T, Mori A, Onodera H, Imamura M (2005) Hepatocyte growth factor as well as vascular endothelial growth factor gene induction effectively promotes liver regeneration after hepatectomy in Solt-Farber rats. Hepatogastroenterology 52: 1393–1397

Oppelt A, Kaschek D, Huppelschoten S, Sison-Young R, Zhang F, Buck-Wiese M, Herrmann F, Malkusch S, Krüger CL, Meub M et al (2018) Model-based identification of TNFα-induced IKKβ-mediated and IκBα-mediated regulation of NFκB signal transduction as a tool to quantify the impact of drug-induced liver injury compounds. NPJ Syst Biol Appl 4: 23

Paranjpe S, Bowen WC, Mars WM, Orr A, Haynes MM, DeFrances MC, Liu S, Tseng GC, Tsagianni A, Michalopoulos GK (2016) Combined systemic elimination of MET and epidermal growth factor receptor signaling completely abolishes liver regeneration and leads to liver decompensation. Hepatology 64: 1711–1724

Puri P, Baillie RA, Wiest MM, Mirshahi F, Choudhury J, Cheung O, Sargeant C, Contos MJ, Sanyal AJ (2007) A lipidomic analysis of nonalcoholic fatty liver disease. Hepatology 46: 1081–1090

Raue A, Kreutz C, Maiwald T, Bachmann J, Schilling M, Klingmuller U, Timmer J (2009) Structural and practical identifiability analysis of partially observed dynamical models by exploiting the profile likelihood. Bioinformatics 25: 1923–1929

Raue A, Schilling M, Bachmann J, Matteson A, Schelker M, Kaschek D, Hug S, Kreutz C, Harms BD, Theis FJ et al (2013) Lessons learned from quantitative dynamical modeling in systems biology. PLoS One 8: e74335

Riazi K, Azhari H, Charette JH, Underwood FE, King JA, Afshar EE, Swain MG, Congly SE, Kaplan GG, Shaheen AA (2022) The prevalence and incidence of NAFLD worldwide: a systematic review and meta-analysis. Lancet Gastroenterol Hepatol 7: 851–861

Ritchie ME, Phipson B, Wu D, Hu Y, Law CW, Shi W, Smyth GK (2015) limma powers differential expression analyses for RNA-sequencing and microarray studies. Nucleic Acids Res 43: e47

Rosenblatt M, Timmer J, Kaschek D (2016) Customized Steady-State Constraints for Parameter Estimation in Non-Linear Ordinary Differential Equation Models. Front Cell Dev Biol 4: 41

Sabapathy T, Helmerhorst E, Ellison G, Bridgeman SC, Mamotte CD (2022) High-fat diet induced alterations in plasma membrane cholesterol content impairs insulin receptor binding and signalling in mouse liver but is ameliorated by atorvastatin. Biochim Biophys Acta Mol Basis Dis 1868: 166372

Schilling M, Maiwald T, Bohl S, Kollmann M, Kreutz C, Timmer J, Klingmuller U (2005) Computational processing and error reduction strategies for standardized quantitative data in biological networks. FEBS J 272: 6400–6411

Schmiester L, Schalte Y, Bergmann FT, Camba T, Dudkin E, Egert J, Frohlich F, Fuhrmann L, Hauber AL, Kemmer S et al (2021) PEtab-Interoperable specification of parameter estimation problems in systems biology. PLoS Comput Biol 17: e1008646

Schwarz G (1978) Estimating the Dimension of a Model. Ann Statist 6: 461–464, 464

Seo J, Jeong DW, Park JW, Lee KW, Fukuda J, Chun YS (2020) Fatty-acid-induced FABP5/HIF-1 reprograms lipid metabolism and enhances the proliferation of liver cancer cells. Commun Biol 3: 638

Slankamenac K, Graf R, Barkun J, Puhan MA, Clavien PA (2013) The comprehensive complication index: a novel continuous scale to measure surgical morbidity. Ann Surg 258: 1–7

Tajima T, Goda N, Fujiki N, Hishiki T, Nishiyama Y, Senoo-Matsuda N, Shimazu M, Soga T, Yoshimura Y, Johnson RS et al (2009) HIF-1α is necessary to support gluconeogenesis during liver regeneration. Biochemical and Biophysical Research Communications 387: 789–794

Tekkesin N, Taga Y, Sav A, Almaata I, Ibrisim D (2011) Induction of HGF and VEGF in hepatic regeneration after hepatotoxin-induced cirrhosis in mice. Hepatogastroenterology 58: 971–979

Tremblay F, Brule S, Hee Um S, Li Y, Masuda K, Roden M, Sun XJ, Krebs M, Polakiewicz RD, Thomas G et al (2007) Identification of IRS-1 Ser-1101 as a target of S6K1 in nutrient- and obesity-induced insulin resistance. Proc Natl Acad Sci U S A 104: 14056–14061

Vivero A, Ruz M, Rivera M, Miranda K, Sacristan C, Espinosa A, Codoceo J, Inostroza J, Vasquez K, Perez A et al (2021) Zinc Supplementation and Strength Exercise in Rats with Type 2 Diabetes: Akt and PTP1B Phosphorylation in Nonalcoholic Fatty Liver. Biol Trace Elem Res 199: 2215–2224

Vizcaino JA, Deutsch EW, Wang R, Csordas A, Reisinger F, Rios D, Dianes JA, Sun Z, Farrah T, Bandeira N et al (2014) ProteomeXchange provides globally coordinated proteomics data submission and dissemination. Nat Biotechnol 32: 223–226

Zhang J, Gao Z, Yin J, Quon MJ, Ye J (2008) S6K directly phosphorylates IRS-1 on Ser-270 to promote insulin resistance in response to TNF-(alpha) signaling through IKK2. J Biol Chem 283: 35375–35382

